# Benchmarking Time-Series Data Discretization on Inference Methods

**DOI:** 10.1101/378620

**Authors:** Yuezhe Li, Tiffany Jann, Paola Vera-Licona

## Abstract

The rapid development in quantitatively measuring DNA, RNA, and protein has generated a great interest in the development of reverse-engineering methods, that is, data-driven approaches to infer the network structure or dynamical model of the system. Many reverse-engineering methods require discrete quantitative data as input, while many experimental data are continuous. Some studies have started to reveal the impact that the choice of data discretization has on the performance of reverse-engineering methods. However, more comprehensive studies are still greatly needed to systematically and quantitatively understand the impact that discretization methods have on inference methods. Furthermore, there is an urgent need for systematic comparative methods that can help select between discretization methods. In this work, we consider 4 published intracellular networks inferred with their respective time-series datasets. We discretized the data using different discretization methods. Across all datasets, changing the data discretization to a more appropriate one improved the reverse-engineering methods’ performance. We observed no universal best discretization method across different time-series datasets. Thus, we propose DiscreeTest, a two-step evaluation metric for ranking discretization methods for time-series data. The underlying assumption of DiscreeTest is that an optimal discretization method should preserve the dynamic patterns observed in the original data across all variables. We used the same datasets and networks to show that DiscreeTest is able to identify an appropriate discretization among several candidate methods. To our knowledge, this is the first time that a method for benchmarking and selecting an appropriate discretization method for time-series data has been proposed.

**Availability:** All the datasets, reverse-engineering methods and source code used in this paper are available in Vera-Licona’s lab Github repository: https://github.com/VeraLiconaResearchGroup/Benchmarking_TSDiscretizations

## 1 Introduction

Understanding important aspects in molecular cell biology requires insight into the structure and dynamics of networks that are made up of thousands of interacting components such as DNA, RNA, proteins and metabolites. One of the central goals of systems biology is to unravel the complex web of interactions among these components (Dasgupta B, 2011). With the rapid development of high-throughput technologies, the amount of biological data that either reports different molecules’ concentration in the form of steady-state or time-series data is constantly increasing. This unprecedented explosion of data has opened the doors to development and improvement of methods that infer or “reverse-engineer” intracellular networks. Several reverse engineering methods infer these networks from discretized time-series data, using diverse modeling frameworks such as dynamic Bayesian networks (Perrin *et al.*, 2003), or Boolean (Liang *et al.*, 1998; Mehra *et al.*, 2004; Martin *et al.*, 2007; Dimitrova *et al.*, 2011; Vera-Licona *et al.*, 2014) and polynomial dynamical systems (Jarrah *et al.*, 2007).

In the most general sense, data discretization is the process of converting continuous features or variables to discrete or nominal features or variables. Discretizing continuous data has been a long-standing problem in data mining and knowledge discovery (see for example, (MacQueen, 1967; Kohonen, 1989; Catlett, 1991)). Different discretization methods have been developed to address different needs: supervised *vs.* unsupervised, dynamic *vs.* static, global *vs.* local, splitting (top-down) *vs.* merging (bottom-up), and direct *vs.* incremental. Examples of unsupervised methods include k-means clustering (MacQueen, 1967), equal width interval (Catlett, 1991; Dougherty *et al.*, 1995; Kerber, 1992a), equal frequency interval (Catlett, 1991; Dougherty *et al.*, 1995; Kerber, 1992a), and graph-theoretic based discretization (Dimitrova *et al.*, 2010). Examples of supervised methods include ChiMerge (Kerber, 1992b), D-2 (Catlett, 1991), Vector Quantization (Kohonen, 1989), Holte’s 1R Discretizer (Holte, 1993) and more recently, for time-series gene expression data, discretization methods like those introduced in (Lustgarten JL, 2011; Misra and Ray, 2017). See (Liu *et al.*, 2002; Kotsiantis S, 2006) for nice surveys on data discretization methods.

In computational systems biology, when inferring intracellular networks from high through-put data, data discretization is an important pre-processing step for many inference (or reverse-engineering) methods that require discrete data as input. Previous studies have reported that by replacing the data discretization methods, the precision and sensitivity of reverse-engineering methods can be significantly improved (Vera-Licona *et al.*, 2014; Li *et al.*, 2010; Velarde *et al.*, 2008).

Our motivation and focus for this work is to systematically examine the impact of different data discretization methods on the performance of reverse-engineering algorithms, with a particular interest in time-series data. We use 4 different sets of published time-series data that have been used to infer, with different reverse-engineering methods, 4 intracellular networks: three gene regulatory networks and one cell signaling network. We first reproduce the results reported by authors in the respective papers by using the same original data, same discretization methods, and same reverse engineering tools. We then alternate through all the discretization methods in GED PRO TOOLS: Gene Expression Data prePROcessing TOOLS (Gallo *et al.*, 2016) to observe the impact of data discretization choice on the performance of reverse engineering methods.

Consistent with what has been observed in previous studies, there is not an optimal data discretization method that works better for all the different datasets. Rather, the problem of data discretization is rather context and data-dependent (Gallo *et al.*, 2016). To address the problem of selecting an optimal discretization method for time-series data, we propose DiscreeTest, a two-step evaluation metric for ranking discretization methods for time-series data. The main assumption behind DiscreeTest’s metric is that the optimal discretization method best preserves the observed dynamic patterns in the original data. We validated the performance of DiscreeTest using the aforementioned published time-series data.

## 2 Materials and Methods

### 2.1 Discretization of time-series data using GED PRO TOOLS

In our study, we use GED PRO TOOLS (Gallo *et al.*, 2016) to discretize the data. GED PRO TOOLS provides 16 different types of discretization methods, 13 of which are unsupervised discretization approaches (Table 1, Table S1). Some of the 16 types of discretization methods discretize the data in multiple levels, such as bikmeans discretization, which can discretize data into 2, 3, 4 or 5 categories. Nine of these discretization methods, such as target discretization threshold (TDT) (Gallo *et al.*, 2011), can only discretize continuous data into 2 categories.

For simplicity, throughout the paper we refer to different discretization methods in our paper not only when distinguishing between different types of discretization method, but also when using the same approach but considering different levels of discretization, such as bikmeans2 (2 categories of discretization) and bikmeans3 (3 categories of discretization).

### 2.2 Intracellular networks, their time-series data and reverse-engineering algorithms used for this study

We use 4 different published intracellular networks with their respective time-series datasets: (1) DREAM3 Yeast In Silico, a yeast (*Saccharomyces cerevisiae*) *in silico* gene regulatory network published in (Marbach *et al.*, 2009, 2010; Prill *et al.*, 2010), (2) Pandapas, an *in silico* gene regulatory network published in (Camacho *et al.*, 2007), (3) IRMA, a yeast synthetic network, published in (Cantone *et al.*, 2009) and, (4) Hepatocytic Cell Signaling network, an *in vivo* signaling network of hepatocellular carcinoma introduced in (Saez-Rodriguez *et al.*, 2009).

Here, we use the same reverse engineering methods to infer each network’s structure as used in its original study.

#### 2.2.1 DREAM3 Yeast In Silico Network

The DREAM3 Yeast In Silico network contains 100 genes and 9900 gene interactions. This network was provided by the DREAM 3 In Silico Network Challenge (Marbach *et al.*, 2010, 2009; Prill *et al.*, 2010). There are 46 different time series. Each time series has 21 time points, and there is a unique perturbation in each time series. The gold standard of this network is known. We examine how data discretization influences the accuracy of time-delayed dynamic Bayesian network (TDBN), originally introduced in (Zou and Conzen, 2005) and later applied in (Li *et al.*, 2014) to infer the DREAM3 yeast *in silico* network. Since TDBN requires binary input data, we select all 11 discretization methods in GED PRO TOOLS capable of binary discretization (bikmeans2, i2, kmeans2, max25, max50, max75, mean, q2, TDT, top25, top75) for comparison.

#### 2.2.2 Pandapas Network

The *in silico* gene regulatory network introduced in (Camacho *et al.*, 2007) contains 13 nodes and 19 interactions; 10 of the nodes represent genes while the other 3 nodes (P1, P2, P3 in Figure S1) represent external perturbations. There are 8 different time series as a result of the different combinations of the 3 perturbation nodes. In (Camacho *et al.*, 2007), BANJO 2.2.0 (Yu *et al.*, 2004) was used to infer the Pandapas network. In addition, we use time-delayed dynamic Bayesian network (Zou and Conzen, 2005) to reverse engineer this network. We test 23 different data discretizations with different levels (bikmeans2, bikmeans3, bikmeans4, bikmeans5, i2, i3, i4, i5, erdals, ji&tan, kmeans2, kmeans3, kmeans4, kmeans5, mean-sd, q2, q3, q4, q5, soinov, TDT, TSD, max50) on inferring the Pandapas network using BANJO, but only the binary discretized data when using TDBN.

#### 2.2.3 IRMA Network

IRMA (*in vivo* “benchmarking” of reverse-engineering and modeling approaches) is a synthetic network that consists of 5 genes (CRF1, GAL4, SWI5, ASH1, and GAL80) and 8 interactions (Figure S3). The network is constructed in a manner such that it can be fully activated with the presence of galactose (switch-on), and inactivated by glucose (switch-off). BANJO 1.0.4, a reverse engineering software (Yu *et al.*, 2004), was utilized in (Cantone *et al.*, 2009) for network inference within the Bayesian network modeling framework. Quantile binary discretization was applied to the data before inputting into BANJO in (Cantone *et al.*, 2009). We use BANJO 1.0.4 to infer network structure using time series (both switch on and off) data. We tested 22 different discretizations with different levels (bikmeans2, bikmeans3, bikmeans4, bikmeans5, i2, i3, i4, i5, erdals, ji&tan, kmeans2, kmeans3, kmeans4, kmeans5, mean-sd, q2, q3, q4, q5, soinov, TDT, TSD).

#### 2.2.4 Hepatocytic Cell Signaling Network

The Hepatocytic Cell Signaling network is a network of 81 nodes and 118 interactions (Figure S4). This network was introduced in (Saez-Rodriguez *et al.*, 2009) to test and validate a reverse engineering method using Cell Network Optimizer (CNO). There are 128 time series collected from CSR (cue-signal-response) from HepG2 hepatocellular carcinoma cells exposed to one of seven cytokines in the presence or absence of seven small-molecule kinase inhibitors. The authors used a genetic algorithm to minimize mean squared errors (MSE) between data generated by the inferred networks and the original experimental data by adding or removing interactions in the prior knowledge network. We notice that size penalty, a parameter that is embedded in the genetic algorithm of CNO, was a constant when (Saez-Rodriguez *et al.*, 2009) was published, but now is a tunable in the current R package (MacNamara A, 2012). We also explore how size penalty range influences network optimization. We discretize our hepatocytic cell signaling data with all the 11 different binary discretization methods in GED PRO TOOLS (bikmeans2, i2, kmeans2, max25, max50, max75, mean, q2, TDT, top25, top75).

**Table 1.** Discretizations used in this study

| abbreviation | full name | levels | technique category |
| --- | --- | --- | --- |
| bikmeansX | bidirectional kmeans discretization with X levels | $2 \leq X \leq 5$ | clustering |
| erdals | Erdal's et.al method (Erdal *et al.*, 2004; Madeira and Oliveira, 2005) | 2 | variation between time points |
| TDT | taget discretization threshold (Gallo *et al.*, 2011) | 2 | metric cutoffs |
| iX | equal width discretization with X levels(Madeira and Oliveira, 2005) | $2 \leq X \leq 5$ | metric cutoffs |
| qX | equal frequency discretization with X levels (Madeira and Oliveira, 2005) | $2 \leq X \leq 5$ | metric cutoffs |
| mean | discretization through comparing to mean value (Madeira and Oliveira, 2005) | 2 | metric cutoffs |
| kmeansX | kmeans discretization with X levels (Li *et al.*, 2010) | $2 \leq X \leq 5$ | clustering |
| ji&tan | Ji and Tan's method (Ji and Tan, 2004) | 3 | variation between time points |
| soinov | Soinov's change state method(Soinov *et al.*, 2003) | 2 | variation between time points |
| mean-sd | mean plus standard (Ponzoni *et al.*, 2007) | 3 | metric cutoffs |
| TSD | translational state discretization (Möller-Levet *et al.*, 2003) | 2 | variation between time points |
| maxY | Max - Y% Max (Madeira and Oliveira, 2005) | 2 | metric cutoffs |
| topY | Top%Y discretization (Madeira and Oliveira, 2005) | 2 | metric cutoffs |

### 2.3 DiscreeTest: A Two-step Benchmark Metric of Time Series Data Discretization Methods

We propose a two-step discretization evaluation metric (DiscreeTest) to benchmark and identify an optimal discretization method for a given time-series dataset. We propose our metric under the assumption that we want the discretized data to preserve dynamic patterns observed in the original data. To that end, DiscreeTest benchmarks, filters and ranks different discretization methods to find the optimal one. Each discretized dataset is subject to two steps: (1) qualification and (2) evaluation (Algorithm 1). For both steps, we normalize both original data and discretized data to have values between 0 and 1.

The qualification step uses sign test to measure whether original data and discretized data have dynamic patterns that are statistically different. We obtain the difference between these two groups of data. Since this difference is from both noise and discretization, we cannot assume any distribution on the difference, but we expect them to separate evenly on the left and right of 0. We then calculate the p-value for running the difference data for a sign test. We then compare this p-value to *α*, the critical value for rejecting the null hypothesis in sign test. If the p-value is larger than *α*, then we fail to reject the null hypothesis and move towards our evaluation step. Otherwise, we disqualify the corresponding discretization method by assigning it an invalid evaluation value (infinite). This qualification step aims to prevent over-fitting the original time series data by adding extra levels/categories in the discretization. In the DiscreeTest Algorithm section in our Supplementary Materials, we provide a detailed example to show how the normalization and sign test work in DiscreeTest).

For the methods that pass the qualification step, the second step in DiscreeTest is to quantitatively evaluate the similarity between dynamic patterns of original data and discretized data. We plot the time points of the normalized original time series and the normalized discretized data; then we interpolate the piecewise-linear curve for our two time series and calculate the mean area between the curves (MABC) of the original data and discretized data, the evaluation value that we return. If the original data and discretized data match perfectly, the mean area between these two curves would be 0. We can identify the best discretization by finding the discretization with the lowest DiscreeTest value. Methods that do not qualify will be ranked last, due to their infinity value assignment. In summary, we consider the optimal data discretization to best keep the intrinsic dynamical trend of the original time series data without over-fitting.

~~~
**Data:** The original data *data_o_* and discretized data *data_d_*.
**Result:** An evaluation of *data_d_*.
Calculate the residue between *data_o_* and *data_d_*
Sign test whether residues of each node in the network has a
median of 0 (*α* = 0.01)
**if** *sign test fails to reject null hypothesis for all nodes* **then**
   return ∞
**end**
*mabc* = mean area between the curves of *data_o_* and *data_d_*
**return** *mabc*;
      **Algorithm 1:** DiscreeTestmetric(*data_o_*, *data_d_*)
~~~

~~~
**Data:** The original data *data_o_* and a list several candidate
      discretization methods *M*.
**Result:** The optimal discretization method.
*minVal* = ∞
*minMethod* = *None*
**for** *method in M* **do**
   *val* = DiscreeTestmetric(*data_o_*, method(*data_o_*))
   **if** *val* < *minVal* **then**
      *minVal* = *val*
      *minMethod* = *method*
   **end**
**end**
**return** *minMethod*;
      **Algorithm 2**: DiscreeTestprocedure(*data_o_*, *M*)
~~~

## 3 Results

### 3.0.1 DREAM3 Yeast In Silico

To begin our experiment, we first find an appropriate value for the maximum time delay, a parameter in the time-delayed dynamic Bayesian network (TDBN) method. We show in Table S2 that 4 is adequate, as a value larger than 4 increases computation complexity without improving performance.

We then use different data discretization methods from GED PRO TOOLS. In (Li *et al.*, 2014; Zou and Conzen, 2005) the authors used the equivalent of the *mean* discretization method in GED PRO TOOLS (threshold by the mean of each variable) before inferring the network using TDBN.

We compute the area under the receiver operating characteristic curves (AUROC) for each reverse-engineered network inferred from differently discretized data. As seen in Figure 1 AUROC for mean discretization is slightly higher than what was reported in (Li *et al.*, 2014) (by 0.0475), but our area under the precision-recall curve (0.0118) is almost the same with the value (0.0155) reported in (Li *et al.*, 2014).

We observed that Top75 is the data discretization that makes TDBN perform best. TDT and max75 also improve TDBN performance, compared to when using mean discretization, the discretization utilized in (Li *et al.*, 2014; Zou and Conzen, 2005). TDBN with input from Top75 discretization shows an increase of 1.6% AUROC compared to TDBN fed with binary data obtained from mean discretization, which is utilized in (Li *et al.*, 2014). Further analysis shows that the higher AUROC of top75 is due to the inference of fewer false positives compared to mean discretization (Table S3). Using data discretized by top75, TDBN correctly identifies 30 edges that otherwise were missing using mean discretized data and reduces 374 false positive edges that are reported by mean discretization.

**Fig. 1:**
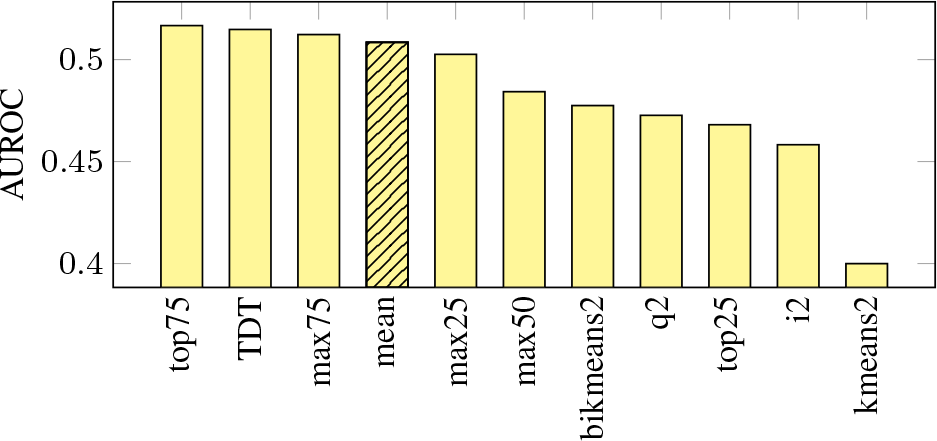
AUROC for the Inference of DREAM3 Yeast In Silico Network Using TDBN Under Different Data Discretizations. Bar plot of the Area Under the ROC Curves (AUROCs) using TDBN with time-series discretized by 11 different binary data discretizations. On the x-axis the 11 discretizations are ordered from left to right to show from the most to the least optimal discretization, according to AUROC. The original publication of TDBN (Li *et al.*, 2014) used Mean discretization as indicated by bar with black stripes. Top75 is the data discretization that makes TDBN method perform best. TDT and max75 also improve TDBN performance, compared to performance when using Mean discretization

### 3.0.2 Pandapas Network

We use both BANJO (version 2.2.0) and time-delayed dynamic Bayesian network (TDBN) to infer Pandapas network structure. In (Camacho *et al.*, 2007), equal frequency with 5 levels (equivalent to quantile 5 discretization, q5) discretization was used for the Pandapas data. We present ROC curves of both reverse engineering methods in Figure S2 and their area under the ROC curves (AUROCs) in Figure 2. We can see that i2 is the best discretization for both inference methods.

**Fig. 2:**
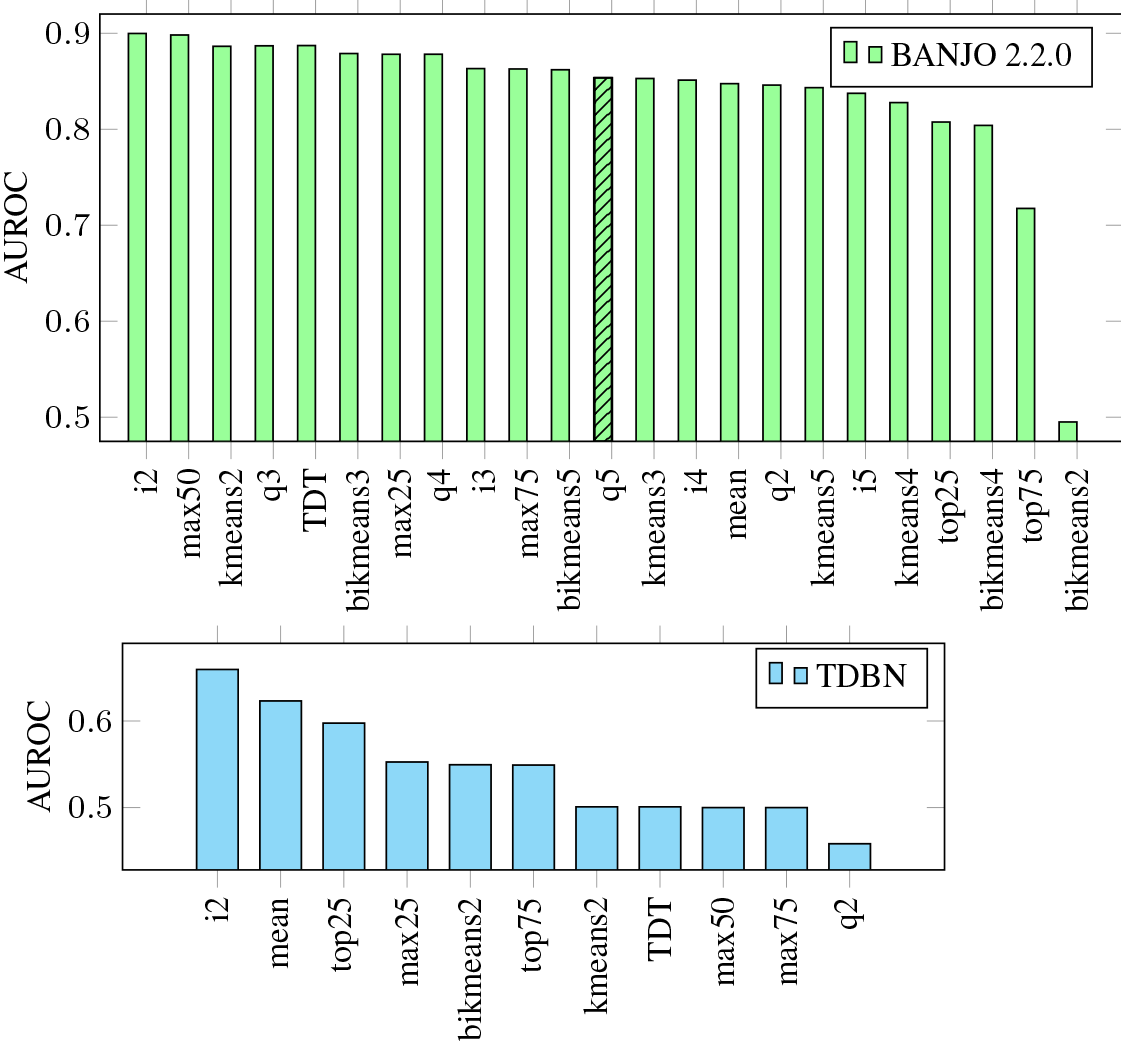
AUROC for the Inference of Pandapas Network Using BANJO 2.2.0 and TDBN Under Different Data Discretizations. Bar plots of the Area Under the ROC Curves (AUROCs) using BANJO 2.2.0 (bar plot in green) and TDBN (bar plot in blue) with time-series discretized by different discretization methods. On the x-axis, the different discretizations are ordered from left to right by most to least optimal, according to AUROC. The original publication inferring Pandapas network with BANJO 2.2.0 (Camacho *et al.*, 2007) used q5 discretization as indicated by bar with black stripes. We can see that i2 is the best discretization for both inference methods.

### 3.0.3 IRMA network

Our experiments with IRMA network using time series with perturbation (switch on and switch off, Figure 3) show that properly choosing a discretization method can largely boost the performance of BANJO, as previously observed in (Vera-Licona *et al.*, 2014). Compared to q2 discretization, which is the discretization method that authors originally used in (Cantone *et al.*, 2009) when considering Switch Off time series, positive predictive value (PPV) can be improved more than 40%, while sensitivity could improve more than 65% when q2 is replaced by q3. In fact, several discretization methods outperformed q2. Among them, q3 and bikmeans2 are both the most sensitive (0.63). While erdals, soinov, and ji&tan give the highest precision (1), they are low in sensitivity (0.13). Mean discretization gives neither the highest sensitivity (0.5) nor positive predicted value (0.8), but its sensitivity is 3 times higher than those have highest positive predicted value, while its positive predicted value is 60% higher than the discretization that gives the highest sensitivity. In summary, for Switch Off time-series data, among the discretization methods tested, there is no best data discretization.

For Switch On time series, TDT, mean, mean-sd and i3 discretizations present better reverse engineering results than q2 discretization, both higher in positive predictive values (PPVs) and sensitivities. TDT discretization gives the highest PPV (0.71), while mean discretization gives the highest sensitivity (0.75).

**Fig. 3:**
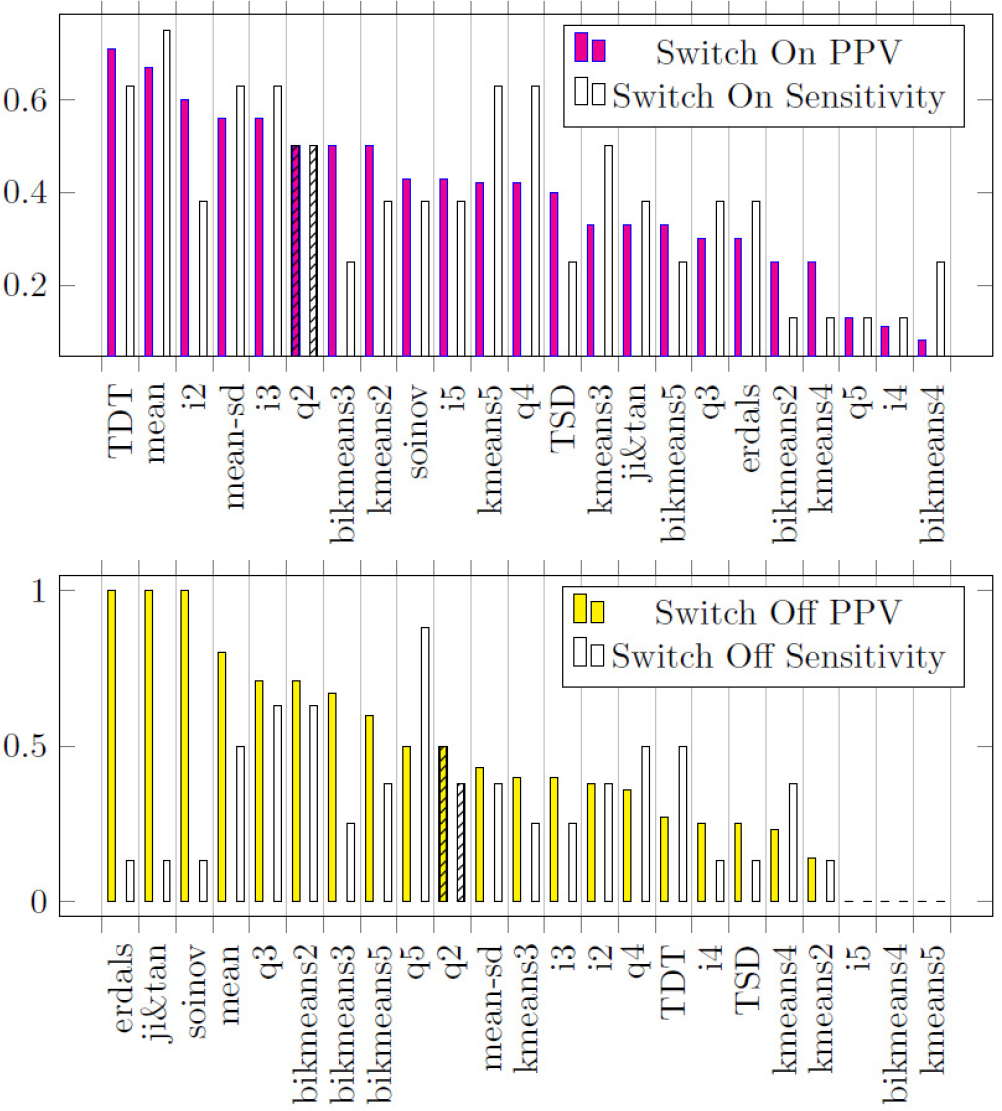
PPV and Sensitivity for the Inference of IRMA network using BANJO 1.0.4 under different data discretizations. Bar plots of Positive Predictive Value (PPV) and Sensitivity for the Inference of IRMA network using BANJO 1.0.4 under different data discretizations. First bar plot, in pink, corresponds to results using Switch On time-series. Second bar plot, in yellow, corresponds to results using Switch Off time-series. On the x-axis the different discretizations. The original publication inferring IRMA network with BANJO 1.0.4 (Cantone *et al.*, 2009) used q2 to discretize both Switch On and Switch Off time-series as indicated by bar with black stripes in both plots.

### 3.0.4 Hepatocytic Cell Signaling Network

We consider the 11 binary data discretizations in GED PRO TOOLS and input the different discretized sets to CNO with different size penalty ranges (Table 2). With the default parameters, we were able to produce the same results reported in (Saez-Rodriguez *et al.*, 2009) (first column of Table 2). We notice that size penalty has a significant impact on optimizing cell signaling network scores. As the size penalty increases, the optimized networks become sparser and eventually, with no edges left. When size penalty is small enough, the variation across discretization methods starts to impact MSE score; for example, bikmeans2 gives universally smaller MSE scores, which is consistent with (Li *et al.*, 2010). However, it is also worth noticing that the network structures with optimal MSE scores are not necessarily the same even if their MSE score is the same: consider the optimized CSR networks with (Lower Bound, Upper Bound) = (0.1, 10) with data discretized by bikmeans2, mean and max50 as shown in Figures S8, S9, S10.

### 3.0.5 Validation of DiscreeTest

From our previous subsections, we have prior knowledge about the structure of the networks that allows us to evaluate the performance of the corresponding reverse-engineering algorithm of choice. For a real application, however, how do we select an optimal discretization method among several methods available? To that end, we propose DiscreeTest, a two-step discretization evaluation method to benchmark and identify an optimal discretization method. DiscreeTest aims to identify the discretization that best retains the dynamic pattern of the original data without over-fitting. We validate DiscreeTest on all our four networks.

#### Validation of DiscreeTest with DREAM3 Yeast In Silico network

In the qualification step, only top75 (p-values > 0.078) and q2 (p-values > 0.039) do not fail. In the evaluation step, top75 provides a smaller mean area between the curves (0.3558) than q2 (0.3676). According to DiscreeTest, top75 is considered to be the best discretization method, which is supported by our previous computations, in which top75 gives the maximum area under the ROC curve using the time-delayed dynamic Bayesian network (TDBN). Here, DiscreeTest identifies the best discretization we found in our reverse engineering experiments.

#### Validation with Pandapas network

In the qualification step, 4 discretization methods pass: top25 (p-value > 0.507), top75 (p-value > 0.179), i2 (p-value > 0.039), mean (p-value > 0.039). Amongst them, i2 gives the minimal mean area between the curves (0.358). This is consistent with the observation that both BANJO and TDBN give the maximum AUROC when using i2 discretized data for network structure inference. Therefore, we conclude DiscreeTest is adequate in this case.

#### Validation with IRMA network

For Switch Off time series, only q2 (p-values>0.99) and q3 (p-values>0.18) pass the qualification step. Among these two discretization methods, q3 gives the smaller MABC (0.4623) (Figure S7). It is also observed that q3 gives a high positive predicted value (0.71) and sensitivity (0.63). Therefore, we conclude DiscreeTest is adequate in this case.

**Table 2.** Mean Squared Error (MSE) from Cell Network Optimizer with Different Size Penalty Range for Hepatocytic Cell Signaling Network

| Lower Bound of Size Penalty | 0.1 | 0.01 | 0.001 | 0.01 | 0.001 | 1 |
| --- | --- | --- | --- | --- | --- | --- |
| Upper Bound of Size Penalty | 10 | 10 | 0.1 | 0.1 | 0.01 | 100.1 |
| bikmeans2 | 0.184 | 0.0018 | $1.010 \times 10^{-5}$ | 0.0041 | $1.837 \times 10^{-5}$ | 16.930 |
| mean | 0.184 | 0.0018 | $1.836 \times 10^{-5}$ | 0.0055 | $1.837 \times 10^{-5}$ | 18.375 |
| q2 | 0.184 | 0.0018 | $1.836 \times 10^{-5}$ | 0.0058 | $1.837 \times 10^{-5}$ | 14.219 |
| max25 | 0.184 | 0.0018 | $1.836 \times 10^{-5}$ | 0.0158 | $1.978 \times 10^{-5}$ | 17.477 |
| top75 | 0.184 | 0.0018 | $1.836 \times 10^{-5}$ | 0.0077 | $1.837 \times 10^{-5}$ | 19.450 |
| i2 | 0.184 | 0.0018 | $1.968 \times 10^{-5}$ | 0.0018 | $2.258 \times 10^{-5}$ | 13.846 |
| max75 | 0.184 | 0.0018 | $2.258 \times 10^{-5}$ | 0.0018 | $1.837 \times 10^{-5}$ | 18.371 |
| TDT | 0.189 | 0.0018 | $3.938 \times 10^{-5}$ | 0.0018 | $1.837 \times 10^{-5}$ | 12.481 |
| kmeans2 | 0.184 | 0.0018 | $3.938 \times 10^{-5}$ | 0.0077 | $1.837 \times 10^{-5}$ | 19.259 |
| max50 | 0.191 | 0.0099 | 0.0261 | 0.216 | 0.0171 | 17.848 |
| top25 | 0.187 | 0.0018 | $1.836 \times 10^{-5}$ | 0.0058 | $1.940 \times 10^{-5}$ | 18.467 |

For Switch On time series, mean (p-values>0.02), q2 (p-values>0.8), q3 (p-values>0.2), q4 (p-values>0.07), and q5 (p-values>0.02) discretizations pass the qualification step, with mean discretization yielding the lowest mean area between the curves (0.374) (Figure S7). However, for this case, we do not observe a discretization that gives a universal good result, i.e. both high sensitivity and high positive predicted value. Mean discretization, nevertheless, gives a high sensitivity (0.75) and a high positive predicted value (0.8)–the second highest positive predicted value among all discretization methods. Therefore, we conclude DiscreeTest is adequate in this case.

#### Validation with Hepatocytic Cell Signaling network

There are only 3 discretization methods that pass the qualification step: bikmeans2(p-value = 0.059), kmeans2 (p-value = 0.25), and top75 (p-value = 0.043). Among them, bikmeans2 gives lowest mean area between the curves value (0.137), as well as the lowest MSE score (0.135). Therefore, DiscreeTest’s identified optimal discretization method is the one that makes CNO perform best.

#### Assessment of Impact of Noise in the Data on DiscreeTest

Each discretization and each reverse-engineering method processes noise in different ways. Furthermore, when combined, the discretization and the reverse engineering methods will process the noise in an intertwined manner. To provide an overview on the impact of noise on our evaluation metric, we assess the performance of DiscreeTest in two ways:

1. **Assessment of noise impact after discretization and reverse-engineering methods have been applied:** In the paragraphs above, we assess DiscreeTest using both *in silico* and real networks. The time-series from both the IRMA network and the Hepatocytic Cell Signaling (CSR) Network are experimental, thus intrinsically contain noise. In both cases, we show that DiscreeTest chooses an optimal discretization method for these network and their timeseries datasets.
2. **Assessment of noise impact before reverse-engineering methods are been applied:** The objective is to assess whether DiscreeTest can identify an optimal discretization method that captures dynamic trends observed in the data, with different levels of noise. We considered the *in silico* Pandapas network and its corresponding time-series datasets. We add 0%, 1% and 5% noise to the time-series datasets before discretizing. In the “Assessment of Data-noise Impact on DiscreeTest” section in the Supplementary Materials, we compared the dynamic patterns in the original time series and the time-series under several discretization methods. We show that the discretization methods identified by DiscreeTest consistently give higher positive predictive value and high accuracy across noise levels (see Tables S8-S10). Furthermore, dynamic patterns observed in the data are shown to be better captured by data discretized by methods identified as optimal by DiscreeTest under the different levels of noise considered (Figure S10).

## 4 Discussion

In this paper, we show that data discretization can have a strong impact on the performance of reverse engineering algorithms. We discuss a wide range of data discretization methods, some of which have multiple levels. Our experiments on 4 different networks inferred by 3 different reverse engineering algorithms reveal no universally optimal data discretization, either in method or in discretization level: For the Hepatocytic Cell Signaling network inference with cell network optimizer (CNO), we observe that bikmeans2 is one of the best choices to discretize the input time-series data, which is consistent with (Li *et al.*, 2010). For the Yeast in Silico network inference using time-delayed dynamic Bayesian network (TDBN), top75 is the best choice for data discretization. For the Pandapas network inference with both BANJO and TDBN, we see that i2 is the data discretization that makes both inference methods perform the best. This observation is notable, as DiscreeTest is able to identify one discretization method that gives two different reverse engineering algorithms best performance. Finally, for the IRMA network inference with BANJO, among the methods tested, there is no best data discretization, since none of them lead BANJO to have both highest positive predicted value and sensitivity for either Switch On or Switch Off time series. However, DiscreeTest identified top candidate discretization methods for both Switch On and Switch Off time-series.

We also notice that for the Hepatocytic Cell Signaling network, even if two inferred networks with CNO have the same MSE score, their corresponding network structures can be different. Comparing the network inferred from bikmeans2 discretized data to the network inferred from Mean discretized data, we observe some new interactions in the network with bikmeans2 discretized data (AKT s→ mTOR, mTOR → IRS1, mTOR → p70S6, p90RSKn → cfos, ras → map3k1) supported by more recent scientific publications (Xia *et al.*, 2014; Yin *et al.*, 2015; Liu *et al.*, 2016; Wan *et al.*, 2016; Gómez-Gómez *et al.*, 2013). There are also some interactions, such as JNK12 → JNK12n, which stand for JNK12 translocation from cytosol to nuclear, that show up in the network inferred using bikmeans2 discretized data. Even though we did not identify any scientific publication supporting this translocation in HepG2 hepatocellular carcinoma cells, this interaction is supported in (Zanella *et al.*, 2008) for human bone osteosarcoma epithelial cells (U2OS Line). These unique interactions suggest that the reverse-engineering network using the data discretized by bikmeans2 is better, not only in score, but also in its capacity to infer biologically meaningful and novel network interactions.

We propose DiscreeTest, a metric with potential to benchmark and identify optimal discretization method(s) independent of the choice of reverse-engineering method. It is essential to remember that the DiscreeTest metric is developed under the assumption that it is desirable for the discretized time series to have a similar dynamic pattern as the original data. Under this assumption, DiscreeTest may not be applied to steady state data. It is worth noticing that the qualification step in DiscreeTest is essential, as it balances between being robust to noise and satisfying the dynamic patterns observed in the original data. A quantitative assessment of the similarity between original data and discretized data is done by calculating the difference between the original data and discretized data over time. Since this difference is from both noise and discretization, we cannot assume any distribution on the difference, but we expect them to separate evenly on the left and right of 0. Thus, we utilize sign test in the qualification step to prevent over-fitting that can arise from the evaluation step, mean area between curves over time (MABC). ABC is the area between the interpolated curve of the original data and the interpolated curve of the discretized data. MABC scales ABC inversely by the number of data points. Therefore, increasing the number of discretization levels can reduce the MABC and in consequence, overfit the data, even if dynamic patterns are different. This explains why some discretization methods yield small MABC, but cannot pass the qualification step. In this context, we showed that DiscreeTest works well in the presence of noise in the data: first, we used the IRMA network and the Hepatocytic Cell Signaling (CSR) network. These datasets inherently contain noise and in both cases we showed that DiscreTest was able to correctly identify an appropriate discretization method. Then, using the *in silico* Pandapas network and its corresponding time series under different levels of noise, we showed that the discretization methods identified by DiscreeTest consistently give higher positive predictive value and high accuracy across different noise levels. Furthermore, the dynamic patterns observed in the original data were shown to be better captured by data discretized by methods identified by DiscreeTest under the different levels of noise considered.

## 5 Conclusion

The choice of a proper data discretization method can largely improve accuracy and sensitivity of reverse engineering algorithms when inferring network structure from discretized time series data. Our experiments show there is not a universally optimal data discretization method. The data discretization method that makes the reverse engineering method perform best depends, at least partially, on the data itself. The two-step discretization evaluation metric (DiscreeTest) is an adequate benchmark and assessment for an optimal discretization method for time-series. The optimality criterion is based on the assumption that an optimal discretization of the data should preserve the dynamic patterns observed in the original data.

## 6 List of abbreviations

MSE: mean squared error; CSR:cue-signal-response; PKN: prior knowledge network; TDBN: time-delayed dynamic Bayesian network; ROC curve: receiver operating characteristic curve; AUC: area under the curve.; AUROC: area under the ROC curve; bikmeansX: bidirectional kmeans discretization with X levels; erdals: Erdal’s et.al discretization method (Erdal *et al.*, 2004); TDT: target discretization threshold; iX: equal width discretization with X levels; qX: equal frequency discretization with X levels; mean: discretization through comparing to mean value; kmeansX: kmeans discretization with X levels; ji&tan: Ji and Tan’s discretization method (Ji and Tan, 2004); soinov: Soinov’s change of state method (Soinov *et al.*, 2003); mean-sd: mean plus standard discretization method; TSD: transilational state discretization; maxY: Max - Y% Max discretization; topY: Top%Y discretization; DiscreeTest: Two-step discretization evaluation.

## 8 Author’s contributions

PVL conceived the project. PVL, YL and TJ designed DiscreeTest. YL performed data discretizations and network analyses. PVL, YL and TJ wrote the manuscript.

## 9 Acknowledgements

TJ worked on this project under the NSF Research Experience for Undergraduates project Modeling and Simulation in Systems Biology (DMS-1460967). We thank Pedro Mendes (University of Connecticut Health) for providing gene expression data for Pandapas network.

## References

Camacho, D., Vera-Licona, P., Mendes, P., and Laubenbacher, R. (2007). Comparison of reverse-engineering methods using an in silico network. Annals of the New York Academy of Sciences, 1115(1), 73–89.

Cantone, I., Marucci, L., Iorio, F., Ricci, M. A., Belcastro, V., Bansal, M., Santini, S., Di Bernardo, M., Di Bernardo, D., and Cosma, M. P. (2009). A yeast synthetic network for in vivo assessment of reverse-engineering and modeling approaches. Cell, 137(1), 172–181.

Catlett, J. (1991). On changing continuous attributes into ordered discrete attributes. In European working session on learning, pages 164–178. Springer.

Dasgupta B, Vera-licona P, S. E. (2011). Reverse engineering of molecular networks from a common combinatorial approach. In Algorithms in Computational Molecular Biology, pages 941–953. John Wiley & Sons, Inc.

Dimitrova, E., Garcia-Puente, L. D., Hinkelmann, F., Jarrah, A. S., Laubenbacher, R., Stigler, B., Stillman, M., and Vera-Licona, P. (2011). Parameter estimation for boolean models of biological networks. Theoretical Computer Science, 412(26), 2816–2826.

Dimitrova, E. S., Licona, M. P. V., McGee, J., and Laubenbacher, R. (2010). Discretization of time series data. Journal of Computational Biology, 17(6), 853–868.

Dougherty, J., Kohavi, R., and Sahami, M. (1995). Supervised and unsupervised discretization of continuous features. In Machine Learning Proceedings 1995, pages 194–202. Elsevier.

Erdal, S., Ozturk, O., Armbruster, D., Ferhatosmanoglu, H., and Ray, W. C. (2004). A time series analysis of microarray data. In Bioinformatics and Bioengineering, 2004. BIBE 2004. Proceedings. Fourth IEEE Symposium on, pages 366–375. IEEE.

Gallo, C. A., Carballido, J. A., and Ponzoni, I. (2011). Discovering time-lagged rules from microarray data using gene profile classifiers. BMC bioinformatics, 12(1), 1.

Gallo, C. A., Cecchini, R. L., Carballido, J. A., Micheletto, S., and Ponzoni, I. (2016). Discretization of gene expression data revised. Briefings in Bioinformatics, 17(5), 758–770.

Gómez-Gómez, Y., Organista-Nava, J., and Gariglio, P. (2013). Deregulation of the mirnas expression in cervical cancer: human papillomavirus implications. BioMed research international, 2013.

Holte, R. (1993). Very simple classification rules perform well on most commonly used datasets. Machine Learning, 11(1), 63–90.

Jarrah, A. S., Laubenbacher, R., Stigler, B., and Stillman, M. (2007). Reverse engineering polynomial dynamical systems. Advances in Applied Mathematics, 39(4), 477–489.

Ji, L. and Tan, K.-L. (2004). Mining gene expression data for positive and negative co-regulated gene clusters. Bioinformatics, 20(16), 2711–2718.

Kerber, R. (1992a). Chimerge: Discretization of numeric attributes. In Proceedings of the Tenth National Conference on Artificial Intelligence, AAAI’92, pages 123–128. AAAI Press.

Kerber, R. (1992b). Chimerge: Discretization of numeric attributes. In Proceedings of the Tenth National Conference on Artificial Intelligence, AAAI’92, pages 123–128. AAAI Press.

Kohonen, T. (1989). Self-organization and Associative Memory: 3rd Edition. Springer-Verlag New York, Inc., New York, NY, USA.

Kotsiantis S, K. D. (2006). Discretization techniques: a recent survey. GESTS Int Trans Comput Sci Eng., 6(4), 393–423.

Li, P., Gong, P., Li, H., Perkins, E. J., Wang, N., and Zhang, C. (2014). Gene regulatory network inference and validation using relative change ratio analysis and time-delayed dynamic bayesian network. EURASIP Journal on Bioinformatics and Systems Biology, 2014(1), 1.

Li, Y., Liu, L., Bai, X., Cai, H., Ji, W., Guo, D., and Zhu, Y. (2010). Comparative study of discretization methods of microarray data for inferring transcriptional regulatory networks. BMC bioinformatics, 11(1), 520.

Liang, S., Fuhrman, S., and Somogyi, R. (1998). Reveal, a general reverse engineering algorithm for inference of genetic network architectures. Pacific Symposium on Biocomputing.

Liu, F., Zhang, W., Yang, F., Feng, T., Zhou, M., Yu, Y., Yu, X., Zhao, W., Yi, F., Tang, W., et al. (2016). Interleukin-6-stimulated progranulin expression contributes to the malignancy of hepatocellular carcinoma cells by activating mtor signaling. Scientific reports, 6.

Liu, H., Hussain, F., Tan, C. L., and Dash, M. (2002). Discretization: An enabling technique. Data Mining and Knowledge Discovery, 6(4), 393–423.

Lustgarten JL, Visweswaran S, G. V. C. G. (2011). Application of an efficient bayesian discretization method to biomedical data. BMC Bioinformatics, 12(309).

MacNamara A (2012). CNORdt: Add-on to CellNOptR: Discretized time treatments.

MacQueen, J. (1967). Some methods for classification and analysis of multivariate observations. In Proceedings of the Fifth Berkeley Symposium on Mathematical Statistics and Probability, Volume 1: Statistics, pages 281–297, Berkeley, Calif. University of California Press.

Madeira, S. C. and Oliveira, A. L. (2005). An evaluation of discretization methods for non-supervised analysis of time-series gene expression data. Instituto de Engenharia de Sistemas e Computadores Investigacao e Desenvolvimento, Technical Report, 42.

Marbach, D., Schaffter, T., Mattiussi, C., and Floreano, D. (2009). Generating realistic in silico gene networks for performance assessment of reverse engineering methods. Journal of computational biology, 16(2), 229–239.

Marbach, D., Prill, R. J., Schaffter, T., Mattiussi, C., Floreano, D., and Stolovitzky, G. (2010). Revealing strengths and weaknesses of methods for gene network inference. Proceedings of the national academy of sciences, 107(14), 6286–6291.

Martin, S., Zhang, Z., Martino, A., and Faulon, J.-L. (2007). Boolean dynamics of genetic regulatory networks inferred from microarray time series data. Bioinformatics, 23(7), 866–874.

Mehra, S., Hu, W.-S., and Karypis, G. (2004). A boolean algorithm for reconstructing the structure of regulatory networks. Metabolic engineering, 6(4), 326–339.

Misra, S. and Ray, S. S. (2017). Finding optimum width of discretization for gene expressions using functional annotations. Computers in Biology and Medicine, 90(Supplement C), 59–67.

Möller-Levet, C. S., Chu, K., and Wolkenhauer, O. (2003). Dna microarray data clustering based on temporal variation: Fcv with tsd preclustering. Applied Bioinformatics, 2(1), 35–45.

Perrin, B.-E., Ralaivola, L., Mazurie, A., Bottani, S., Mallet, J., and d’Alche Buc, F. (2003). Gene networks inference using dynamic bayesian networks. Bioinformatics, 19(suppl 2), ii138–ii148.

Ponzoni, I., Azuaje, F., Augusto, J., and Glass, D. (2007). Inferring adaptive regulation thresholds and association rules from gene expression data through combinatorial optimization learning. IEEE/ACM Transactions on Computational Biology and Bioinformatics, 4(4), 624–634.

Prill, R. J., Marbach, D., Saez-Rodriguez, J., Sorger, P. K., Alexopoulos, L. G., Xue, X., Clarke, N. D., Altan-Bonnet, G., and Stolovitzky, G. (2010). Towards a rigorous assessment of systems biology models: the dream3 challenges. PloS one, 5(2), e9202.

Saez-Rodriguez, J., Alexopoulos, L. G., Epperlein, J., Samaga, R., Lauffenburger, D. A., Klamt, S., and Sorger, P. K. (2009). Discrete logic modelling as a means to link protein signalling networks with functional analysis of mammalian signal transduction. Molecular systems biology, 5(1), 331.

Soinov, L. A., Krestyaninova, M. A., and Brazma, A. (2003). Towards reconstruction of gene networks from expression data by supervised learning. Genome biology, 4(1), 1.

Velarde, C., Rubio-Escudero, C., and Romero-Zaliz, R. (2008). Boolean networks: a study on microarray data discretization. ESTYLF08, Cuencas Mineras (Mieres-Langreo), pages 17–19.

Vera-Licona, P., Jarrah, A., Garcia-Puente, L. D., McGee, J., and Laubenbacher, R. (2014). An algebra-based method for inferring gene regulatory networks. BMC systems biology, 8(1), 1.

Wan, Z. Y., Tian, J. S., Tan, H. W. S., Chow, A. L., Sim, A. Y. L., Ban, K. H. K., and Long, Y. C. (2016). Mechanistic target of rapamycin complex 1 (mtorc1) is an essential mediator of metabolic and mitogenic effects of fgf19 in hepatoma cells. Hepatology.

Xia, J., Guo, S., Fang, T., Feng, D., Zhang, X., Zhang, Q., Liu, J., Liu, B., Li, M., and Zhu, R. (2014). Dihydromyricetin induces autophagy in hepg2 cells involved in inhibition of mtor and regulating its upstream pathways. Food and Chemical Toxicology, 66, 7–13.

Yin, Y., Hua, H., Li, M., Liu, S., Kong, Q., Shao, T., Wang, J., Luo, Y., Wang, Q., Luo, T., et al. (2015). mtorc2 promotes type i insulin-like growth factor receptor and insulin receptor activation through the tyrosine kinase activity of mtor. Cell Research.

Yu, J., Smith, V. A., Wang, P. P., Hartemink, A. J., and Jarvis, E. D. (2004). Advances to bayesian network inference for generating causal networks from observational biological data. Bioinformatics, 20(18), 3594–3603.

Zanella, F., Rosado, A., García, B., Carnero, A., and Link, W. (2008). Chemical genetic analysis of foxo nuclear–cytoplasmic shuttling by using image-based cell screening. Chembiochem, 9(14), 2229–2237.

Zou, M. and Conzen, S. D. (2005). A new dynamic bayesian network (dbn) approach for identifying gene regulatory networks from time course microarray data. Bioinformatics, 21(1), 71–79.

