## Supplementary File for "Benchmarking Time-Series Data Discretization on Inference Methods"

### Supplementary Data

#### Methods and Results

##### Discretizations from GED PRO TOOLS

In Table S1 we show all the discretization methods we use in this study and a brief description of them, including whether they give binary or multi-level discrete data. Among the 13 different types of unsupervised discretization methods provided by GED PRO TOOLS in Gallo et al. [2016], iX is also known as "equal width discretization", as this discretization splits the data range into X equal width intervals, labels them between 0 and $X-1$, and assigns to each observation the label of its corresponding interval. qX is also known as "equal frequency discretization", as the X intervals are drawn so that they have equal number of observations. Each observation is assigned a value equal to the label of its interval (between 0 and $X-1$). Kmeans clustering separates observations into several clusters, and discretization is based on these clusters. It is worth noticing that kmeans discretization in GED PRO TOOLS does not choose its initial cluster centers randomly. Bi-kmeans discretization has a procedure that builds upon kmeans discretization. TopY discretizes data by assigning the bottom $(1-Y\%)$ observations to 0, and the rest to 1. MaxY discretization assigns 0 to all observations smaller than $Y\%$ of the maximum value.

**Table S1:** Discretization methods in GED PRO TOOLS utilized in our study

| abbreviation | full name | levels | calculation |
|---|---|---|---|
| bikmeansX | bidirectional kmeans discretization with X levels Li et al. [2010] | 2-5 | k-means clustering using both gene profiles and column profiles |
| erdals | Erdal's et.al method Erdal et al. [2004], Madeira and Oliveira [2005] | 2 | assign 1 if gene expression level is changes; otherwise 0. |
| TDT | target discretization threshold Gallo et al. [2011] | 2 | $\min(\text{var}(S_1) + \text{var}(S_2))$, where $S_i$ represents a gene state $\text{var}(S_i)$ are variance for $S_i$, $i = 1, 2$, $S_1 \cap S_2 = \emptyset$ |
| iX | equal width discretization with X levelsMadeira and Oliveira [2005] | 2-5 | discretize data by splitting the range of data into X intervals equally |
| qX | equal frequency discretization with X levels Madeira and Oliveira [2005] | 2-5 | split data into strata with each strata having the same amount of data |
| mean | discretization through comparing to mean value Madeira and Oliveira [2005] | 2 | $a_{ij} = 1$ if $a_{ij} > E(A_i)$, otherwise $a_{ij} = 0$ where $A_i = (a_{i1}, a_{i1}, ...)$ |
| kmeansX | kmeans discretization with X levels Li et al. [2010] | 2-5 | assign data into $k$ (k is given ) levels through k-means clustering data into $k$ clusters |
| ji&tan | Ji and Tan's method Ji and Tan [2004] | 3 | compare ratio between consecutive samples |
| soinov | Soinov's change state method Soinov et al. [2003], Ponzoni et al. [2007] | 2 | minimize sum of entropy for up-regulating(1) and down-regulating(0) gene states |
| mean-sd | mean plus standard Ponzoni et al. [2007] deviation discretization | 3 | $a_{ij} = -1$ if $a_{ij} < E(A_i) - \sigma(A_i)$<br>$a_{ij} = 0$ if $E(A_i) - \sigma(A_i) \le a_{ij} \le E(A_i) + \sigma(A_i)$<br>$a_{ij} = 1$ if $a_{ij} > E(A_i) + \sigma(A_i)$, $A_i = (a_{i1}, a_{i1}, ...)$ |
| TSD | transilational state discretization Möller-Levet et al. [2003] | 2 | difference between consecutive samples |
| maxY | Max - Y% Max Madeira and Oliveira [2005] | 2 | $a_{ij} - 1$ if $a_{ij} > \max(A_i) \times (1 - Y\%)$, otherwise 0. |
| topY | Top%Y discretization Madeira and Oliveira [2005] | 2 | the expression values that are in the Y% of the highest values are discretized to 1 and the remaining values to 0. |

##### DREAM3 Yeast In Silico Network

The DREAM3 Yeast In Silico Network is provided by DREAM (Dialogue for Reverse Engineering Assessments and Methods) 3 In Silico Network Challenge (Marbach et al. [2010, 2009], Prill et al. [2010]). We use the time series from the Yeast Network that contains 100 genes (InSilicoSize100-Yeast1). There are 46 time series, each of them has 21 time points. The goal is to infer the internal structure of this gene regulatory network. The DREAM 3 Challenge provides the gold standard of this network. This network was studied by Li et al. [2014] using time-delayed dynamic Bayesian network (TDBN) from Zou and Conzen [2005]. TDBN allows time delay when inferring a gene regulatory network. TDBN requires the input data to be binary. In Li et al. [2014], both area under the receiver operating characteristic curve (AUROC) and area under the precision-recall curve (AUPR) were reported.

**Table S2:** TDBN Area Under the Receive Operating Characteristic Curve (AUROC) with Different Discretization Methods and Maximum Time Delay ,DREAM3 Yeast In Silico Network

| Maximum Delay | AUROC | | |
|---|---|---|---|
| | mean | top75 | i2 |
| 1 | 0.5 | 0.5 | 0.5 |
| 2 | 0.5 | 0.5 | 0.5 |
| 3 | 0.5 | 0.5 | 0.5 |
| 4 | 0.5085 | 0.5176 | 0.4583 |
| 5 | 0.5085 | 0.5176 | 0.4583 |
| 6 | 0.5085 | 0.5176 | 0.4583 |
| 6 | 0.5085 | 0.5176 | 0.4583 |
| 7 | 0.5085 | 0.5176 | 0.4583 |
| 8 | 0.5085 | 0.5176 | 0.4583 |
| 9 | 0.5085 | 0.5176 | 0.4583 |
| 10 | 0.5085 | 0.5176 | 0.4583 |

Table S2 shows that a maximum delay of 4 is adequate, as either before or after that point, the AUROC does not change for any discretization. Table S3 depicts Sensitivity and Specificity values for TDBN method using data discretized by Top75 and Mean.

**Table S3:** Sensitivity, Specificity, True Positive (TP), False Negative (FN), True Negative (TN), and False Positive (FP) values of Time-delayed Dynamic Bayesian Network (TDBN), DREAM3 Yeast In Silico Network

| Discretization | Sensitivity | Specificity | TP | FN | TN | FP |
|---|---|---|---|---|---|---|
| top75 | 0.2892 | 0.7478 | 48 | 118 | 7354 | 2480 |
| mean | 0.2831 | 0.7271 | 47 | 119 | 7150 | 2684 |

##### Pandapas Network

The Pandapas network is an *in silico* network of 13 nodes introduced in Camacho et al. [2007]. In this network, 10 nodes are intrinsic genes (G1 - G10) while other 3 nodes (P1 - P3) represent external perturbations (Figure S1). This network is perturbed by introducing its wild type non-function mutations on G1, G2, ..., G10, respectively. Here, we only focus on the wildtype network. In Camacho et al. [2007], authors benchmarked different reverse-engineering algorithms to infer the network under different levels of noise in the data. We start with noiseless data. Together, there are 8 datasets that are generated with different initial conditions and different perturbations (P1, P2, P3 being either 0, 0.01, 0.05, 0.1, or 0.5). We use both BANJO (version 2.2.0) and time-delayed dynamic Bayesian network (TDBN) to infer the Pandapas network based on 8 datasets with different perturbations. For BANJO, we tested 23 different data discretizations, including some multi-level discretization methods (bikmeans3, bikmeans4, bikmeans5, kmeans3, kmeans4, kmeans5, i3, i4, i5, q3, q4, q5). For a given discretization method, we inferred the networks with BANJO on every one of the 8 dataset separately and considered a consensus network where an edge is considered if it appears more than twice amongst the 8 inference results. For the network inferred using TDBN, we test the 11 binary discretizations available in GED PRO TOOLS.

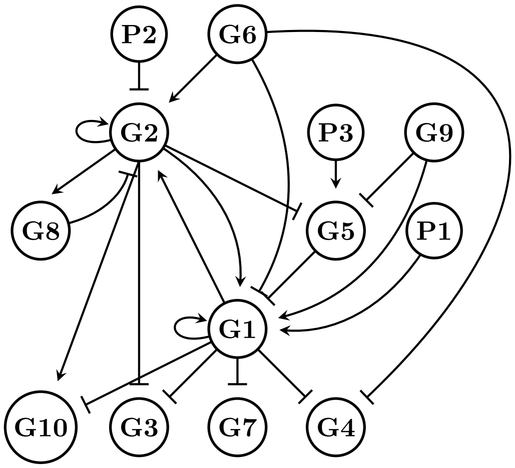


**Figure S1:** Pandapas Network from Camacho et al. [2007]. This is a gene network with 10 genes and 3 environmental perturbations (P1, P2, P3). These perturbations can directly affect the expression rate of gene G1, G2 and G5. Arrow ends mean activation and blunt ends inhibition of the transcription rate.

We present ROC curves of both reverse engineering methods on noiseless data in Figure S2 and their corresponding ROC plots in Fig 2. We can see that i2 is the best data discretization for both BANJO and TDBN.

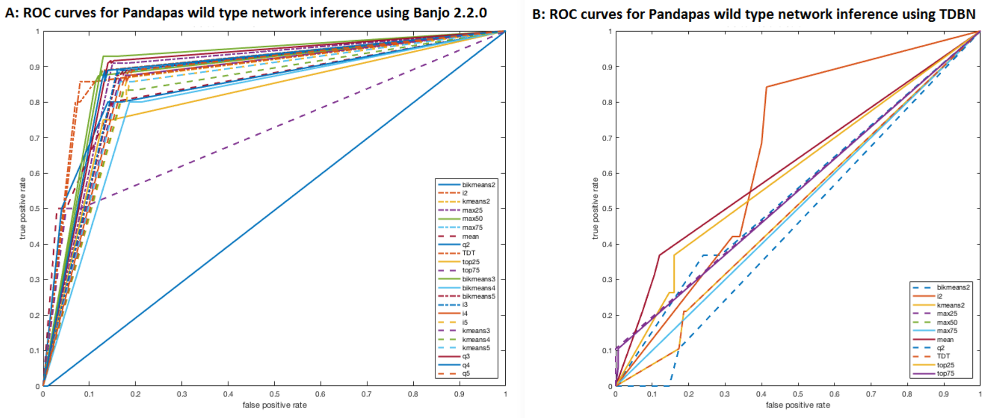


**Figure S2:** ROC Curves for Pandapas Network using either BANJO or TDBN for network inference. A: AUROC of networks inferred by BANJO 2.2.0 using data that is discretized in 23 different ways. Amongst them, i2 gives the maximum AUROC value (0.89983). B: AUROC of networks inferred by TDBN using data that is discretized in 11 different ways. Amongst them, i2 gives the maximum AUROC value (0.659).

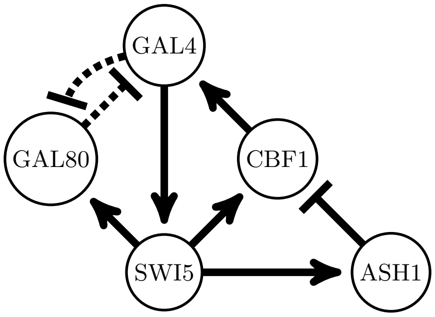


**Figure S3:** IRMA Network from Cantone et al. [2009]. This network consist of 5 different genes: gene GAL80 can be turned off when the environment contains galactose. When galactose is removed from the environment and the yeasts are treated with glucose, the gene expression of IRMA network is turned on. Dashed lines represent inhibitory protein-protein interaction, and directed edges with an arrow end represent activation reactions in the network.

##### IRMA Network

IRMA network is a synthetic yeast network introduced in Cantone et al. [2009]. This network contains 5 genes, CBF1, GAL4, SWI5, ASH1, GAL80 (Figure S3). It is achieved by pairing genes with different promoters and depletion of yeast endogenous transcription factors. Gal80-Gal4 interaction is inhibited in the presence of the glucose. GAL1-10 promoter, cloned upstream of SWI5 gene in the network, is activated by galactose, and consequently, activate all the five network genes. Expression profiles of these genes were analyzed by quantitative real-time RT-PCR (q-PCR). Cantone et al. used BANJO (Yu et al. [2004]) as one of the tools to reverse engineer gene regulatory network from IRMA data. BANJO is a software application and framework for structure learning of static and dynamic Bayesian networks. BANJO focuses on score-based structure inference. In BANJO 1.0.4, there is no effective way to change the cut-off threshold when reporting network. Thereby, we are unable to obtain ROC curves. Thus, we use positive predictive value (PPV) and sensitivity to measure the performance of BANJO when inputting differently discretized time series (switch on and switch off) data.

##### Hepatocytic Cell Signaling Network

The time series data from the Hepatocytic Cell Signaling network was used to infer the network using cell network optimizer (CNO) in Saez-Rodriguez et al. [2009]. Experimental data is collected from HepG2 hepatocellular carcinoma cells. The network is perturbed by exposing one of seven cytokines in the presence or absence of seven small-molecule kinase inhibitors. Before inferring the network structure, all experimental data were normalized, and a prior knowledge network (PKN) was assembled from literature (Terfve et al. [2012], Morris et al. [2011, 2013], Gaudet et al. [2005], Klamt et al. [2006], Saez-Rodriguez et al. [2007, 2008]). This prior knowledge network (PNK), shown in Figure S4, contains edges that are reported in literature in all cell types, thus not all the edges would necessarily exist in HepG2 hepatocellular carcinoma cells. Here, we use their R packages, CellNOptR and CNORdt, released in 2014, for network structure inference (Terfve et al. [2012], Saez-Rodriguez et al. [2009]). Cell network optimizer requires binary discretization, and its default discretization threshold is chosen by the mean value of experimental data after normalizing data using Hill function (Terfve et al. [2012]). The network is optimized through removing edges or changing logical relationships (AND & OR) from the prior knowledge network using a genetic algorithm. Optimized networks are scored through minimizing the mean squared error. In our study, we use Saez-Rodriguez et al. [2009] normalized data, PKN and their default discretization (mean) for comparison . We compare our results of other binary discretizations to mean discretization. We further test how the range of size penalty of cell network optimizer influences the score, which the authors were unable to do but planned to do when Saez-Rodriguez et al. [2009] was published. Using the same parameters in CNO as the authors

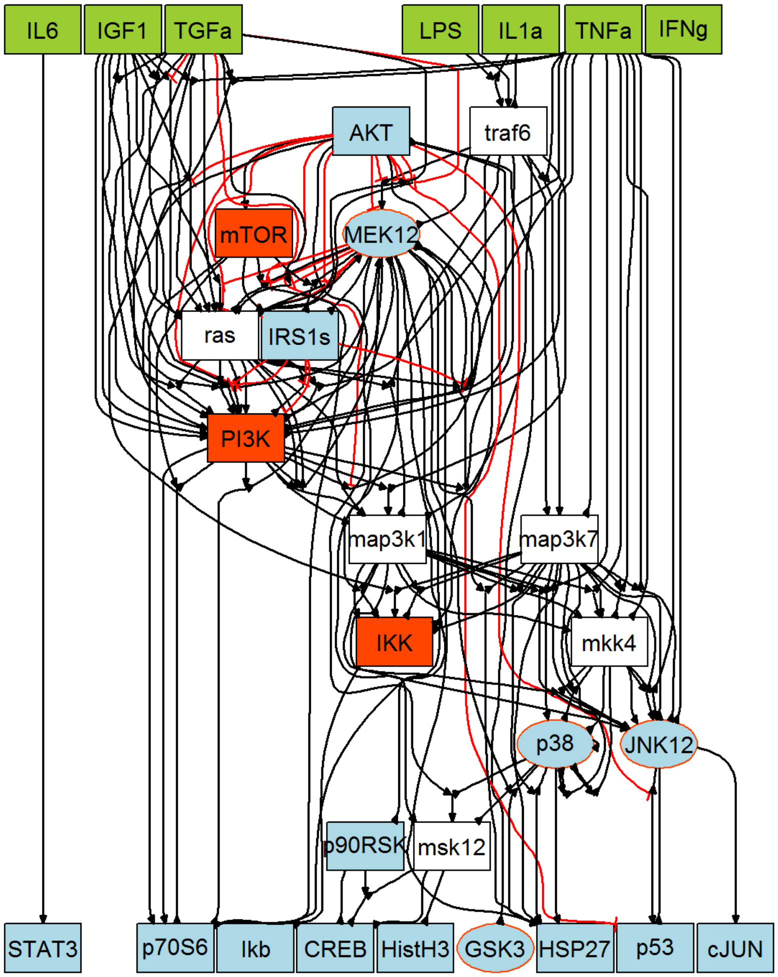


**Figure S4:** The prior knowledge network (PKN) of hepatocytic Cell Signaling network from Saez-Rodriguez et al. [2009]. Green boxes represents 6 cytosines that are tested in this study. They could be either present of absent for each experiment. They influence the function of some proteins directly. Black lines are positive regulatory relations that are reported in literature, red lines are inhibitory regulations from literature.

did in Saez-Rodriguez et al. [2009], bikmeans2 gives a better results than mean; and still does even if some parameters change. Further analysis shows reverse engineered network based on bikmeans2 discretization gives 17 unique edges comparing to the result based on mean discretization. We show detailed networks optimized using data discretized by bikmeans2, mean, and max50 in Figure S5, S6, and S7, respectively.

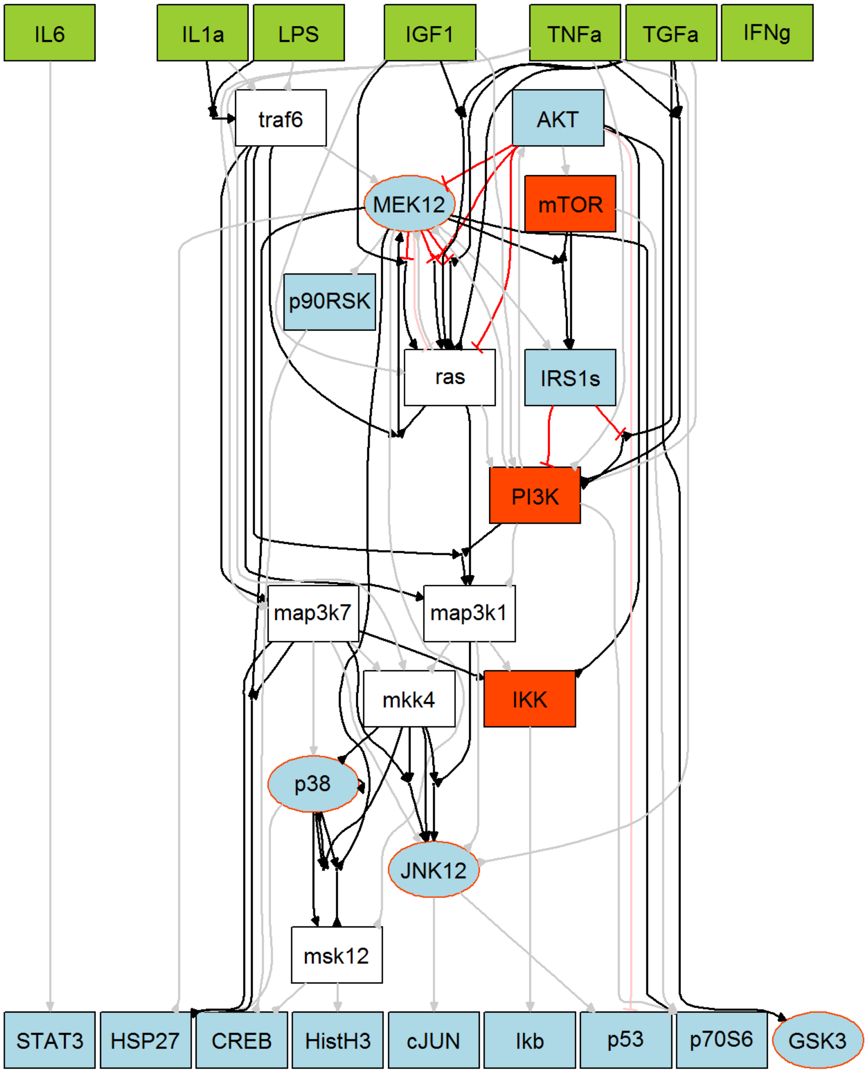


**Figure S5: Optimized CSR Network, (Lower Bound, Upper Bound) = (0.1, 10), Bikmeans2 Discretization.** All the lines shown up are interactions that are reported in literature. Lines with an arrow end represent activation interactions, lines that are red with a blunt end are inhibitory interactions. Grey lines and faded red lines are edges removed during optimization. Black lines are positive regulatory relations, red lines are inhibitory regulations. Green boxes are cytosines. Input data is original data is discretized by bikmeans2 discretization.

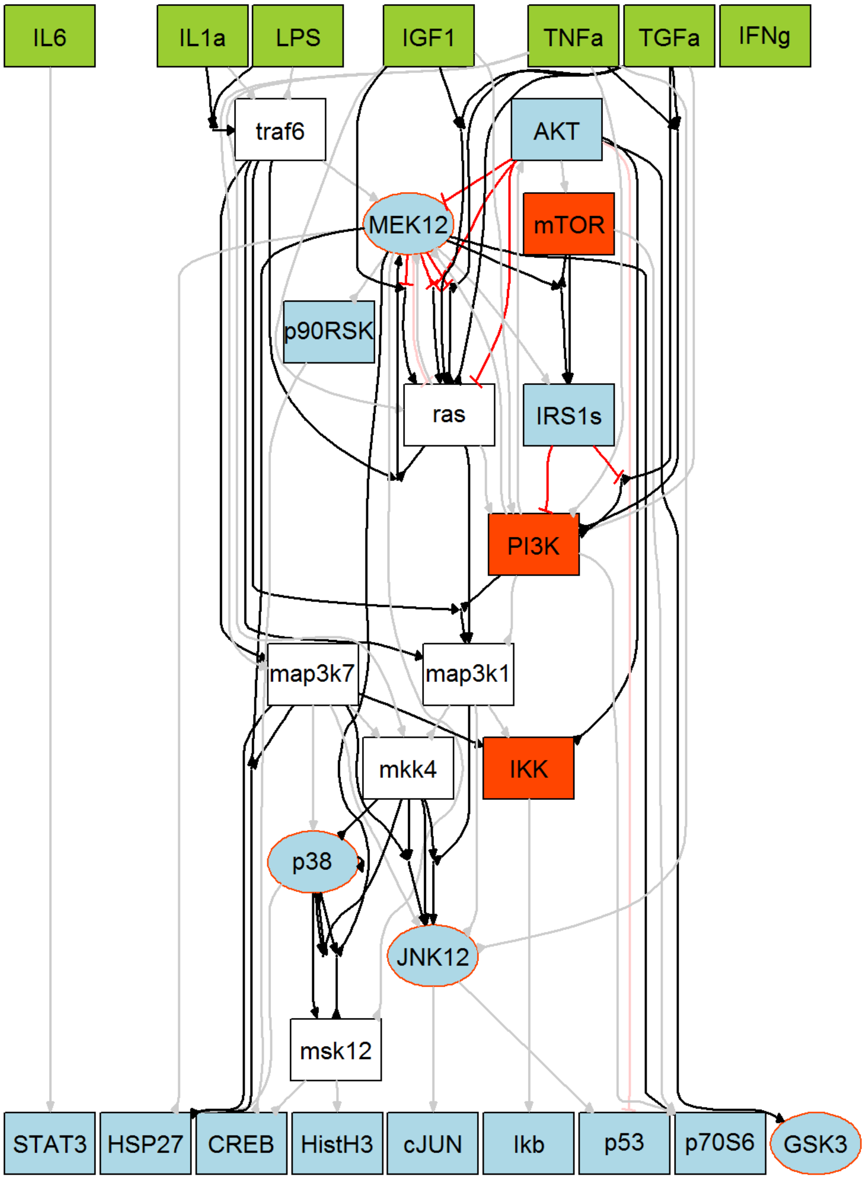


**Figure S6: Optimized CSR Network, (Lower Bound, Upper Bound) = (0.1, 10), Mean Discretization.** All the lines shown up are interactions that are reported in literature. Lines with an arrow end represent activation interactions, lines that are red with a blunt end are inhibitory interactions. Grey lines are edges removed during optimization. Black lines are positive regulatory relations, red lines are inhibitory regulations. Green boxes are cytosines. Input data is original data discretized by mean discretization.

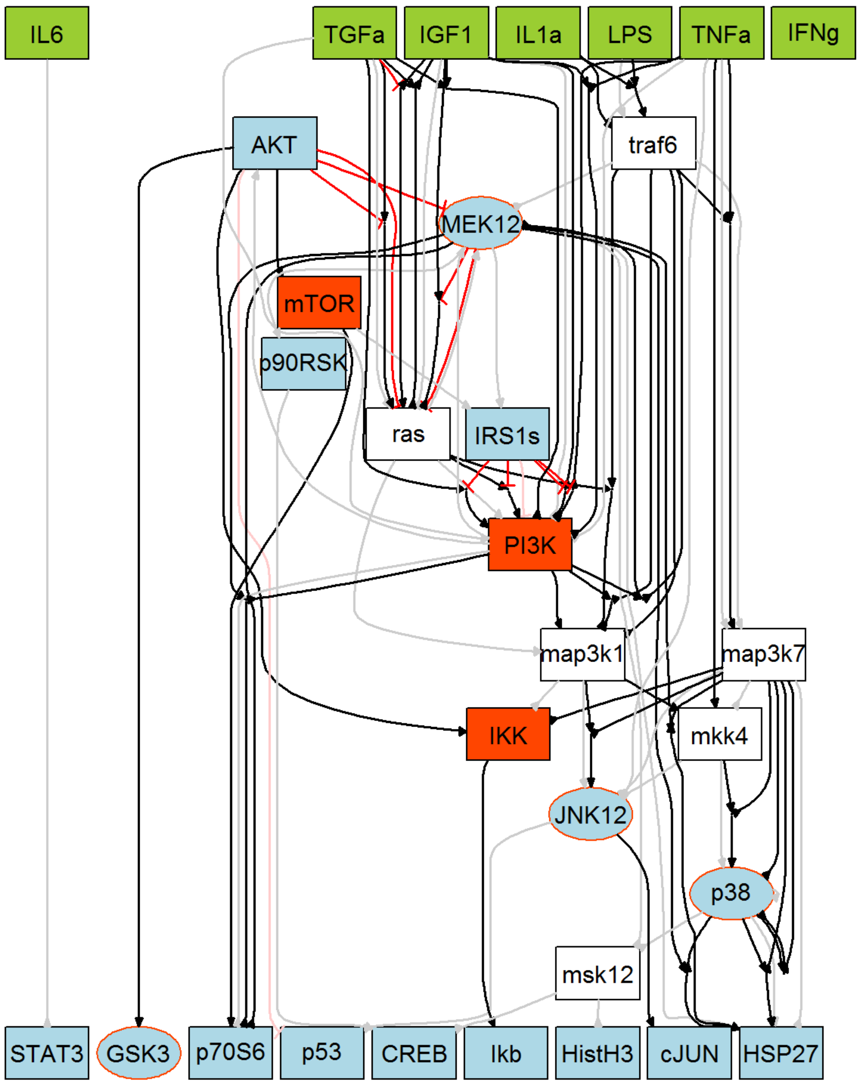


**Figure S7: Optimized CSR Network, (Lower Bound, Upper Bound) = (0.1, 10), Max50 Discretization.** All the lines shown up are interactions that are reported in literature. Lines with an arrow end represent activation interation, lines that are red with a blunt end are inhbitory interactions. Grey lines are edges removed during optimization. Black lines are positive regulatory relations, red lines are inhibitory regulations. Green boxes are cytosines. Input data is original data discretized by max50 discretization.

##### The DiscreeTest Algorithm

For two curves, $y_1 = f(x)$ and $y_2 = g(x)$, $x \in [a, b]$, the mathematical formula for area between the curves (ABC) is defined as $ABC(f, g) = \int_a^b |f(x) - g(x)|dx$. In Figure S8 we can see a graphical representation of the area between curves $f$ and $g$.

For a time series from normalized experimental data,

$$T = \begin{bmatrix} t_{11} & t_{12} & \dots & t_{1m} \\ \vdots & \vdots & \ddots & \vdots \\ t_{n1} & t_{n2} & \dots & t_{nm} \end{bmatrix}$$

where each row is an observation at time point $i$, $1 \le i \le n$ of $m$ different nodes, and $n$ is the total number of time-series datasets. Let $D = (d_{ij})_{n \times m}$ denote the discretized data after normalization (for example, for a discretization with 3 levels, $d_{ij}$ is either 0, 0.5, or 1). We define the mean area between the curves (MA) as

$$MA = \frac{\sum_{j=1}^{m} A(t_j, d_j)}{mn}$$

where $t_j = [t_{1j}, \ldots, t_{nj}]^T$, $d_j = [d_{1j}, \ldots, d_{nj}]^T$, $1 \le j \le m$, $A(t_j, d_j)$ represents the area between the interpolated curves from the normalized original data points and the discretized data points between time points $j-1$ and $j$ (see Figure S9 for an example of the area between the interpolated curves). We also define residue between an original time series after normalization ($t_j$) and discretized and normalized time series ($d_j$) as $r_j$: $r_j = t_j - d_j$. DiscreeTest will used $r_j$ in the first step (qualification step).

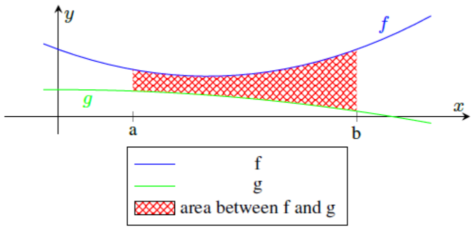


**Figure S8:** An example of the area between the curves $f$ and $g$.

The DiscreeTest algorithm consists of 2 steps: **(1) Qualification Step**. This step consists on testing whether residues of each node ($r_j$, $1 \le j \le m$) reject the null hypothesis in the sign test. When there are multiple time series datasets, DiscreeTest will compute a p-value for each node in each dataset separately. The p-value asisgned to the discretization method will be the lower quantile from all the p-values computed from all the datasets. DiscreeTest uses the lower quantile of all the p-values so that the result is more robust and less likely to be influenced by outliers in the dataset. This p-value will be compared to $\alpha$ (in our study, we choose $\alpha = 0.01$) to determine whether we reject the null hypothesis. Discreetes proceeds when all nodes' residue fail to reject the null hypothesis.

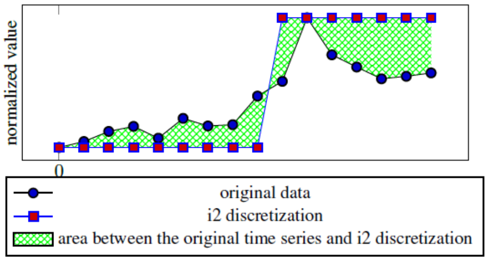


**Figure S9:** An area between the curves example. The original data (black line with blue round dots) is discretized using i2 discretization (blue line with red squared diamond markers). The area highlighted in green is the area between the original time series and i2 discretized time series. Mean area between the curves is an average of total area between the curves over time.

Here we give an example to show how the qualification step works when the discretization method applied has multiple levels. Let's consider gene 3(G3) from the Pandapas Network for which we have 8 time-series datasets. To illustrate how normalization works, let's discretize the data of G3 using the i4 discretization method. The original data for G3 in time-series set 1 is

$$\text{G3}_{\text{orig}} = [0.0329, 0.0445, 0.0619, 0.0674, 0.0676, 0.0666, 0.0657, 0.0651, 0.0648]$$

After applying the i4 discretization method, we have

$$G3_{i4} = [0, 1, 3, 3, 3, 3, 3, 3, 3]$$

The discretized data is normalized to be between 0 and 1 to obtain

$$G3_{i4n} = [0, 0.33, 1, 1, 1, 1, 1, 1, 1]$$

Now, the original data will be normalized to be between 0 and 1 to have now values

$$G3_{n} = [0, 0.3339, 0.8356, 0.9940, 1.0000, 0.9716, 0.9457, 0.9289, 0.9195]$$

Then we compute the residue between the discretized data and the original data is

$$\text{res} = [0, -0.0006, 0.1644, 0.0060, 0, 0.0284, 0.0543, 0.0711, 0.0805]$$

The p-value given by the sign test on this specific sequence of residuals is $p = 0.125$. The same process will be done for each of the 13 nodes of the network and each one of the 8 time-series. Then all the p-values will be pooled together. The lower quartile (25%) of all the 104 p-values is drawn out (p = 0.0039) which is assigned to be the p-value for this discretization. This p-value is smaller than the 0.01, thus this discretization doesnt pass the qualification step, and the evaluation metric of our DiscreeTest is assigned to be infinity.

**(2) Evaluation Step**. The second step is to calculate the mean area between discretized data and original data. We choose the discretization that gives the minimum area between the curves.

In the following tables S4-S7, we report all the p-values and the mean area between the curves (MABC) in all the experiments in the manuscript to provide the reader with a better picture on the performance of the discretization methods tested.

**Table S4:** P-values and Mean Area Between the Curves (MABC) values of Different Discretization for DiscreeTest, DREAM3 Yeast In Silico Network

| Discretization | p-values | MABC |
|---|---|---|
| top75 | 0.078 | 0.356 |
| q2 | 0.022 | 0.368 |
| TDT | p<0.001 | 0.333 |
| max75 | p<0.001 | 0.437 |
| mean | 0.001 | 0.337 |
| max25 | p<0.001 | 0.282 |
| max50 | p<0.001 | 0.360 |
| bikmeans2 | p<0.001 | 0.283 |
| top25 | p<0.001 | 0.426 |
| i2 | p<0.001 | 0.323 |
| kmeans2 | p<0.001 | 0.326 |

**Assessment of data-noise impact on DiscreeTest**.

Each discretization method and each reverse engineering method processes noise in different ways. Furthermore both, the discretization and the reverse engineering methods will process the noise in an intertwined manner. Therefore, it is hard to systematically evaluate how noise impacts our evaluation metric after the discretization and reverse engineering methods have been applied. Thus to provide an overview on the impact of noise on our evaluation metric, we proceed in two ways:

**1. Assessment of noise impact after discretization and reverse-engineering methods have been applied:** In the manuscript we assessed DiscreeTest in both *in silico* and real networks. The time-series from both the IRMA network and the Hepatocytic Cell Signaling (CSR) Network are experimental data

**Table S5:** P-values and Mean Area Between the Curves (MABC) values of Different Discretization for DiscreeTest, Pandapas Network

| Discretization | p-values | MABC |
|---|---|---|
| top25 | 0.5078 | 0.4039 |
| top75 | 0.1797 | 0.4901 |
| i2 | 0.0391 | 0.3581 |
| mean | 0.0391 | 0.3631 |
| bikmeans2 | 0.0039 | 0.2818 |
| TDT | 0.0039 | 0.41 |
| kmeans2 | 0.0039 | 0.41 |
| bikmeans5 | 0.0039 | 0.2601 |
| i4 | 0.0039 | 0.387 |
| q2 | 0.0039 | 0.4281 |
| i5 | 0.0039 | 0.3819 |
| bikmeans4 | 0.0039 | 0.2387 |
| i3 | 0.0039 | 0.3881 |
| kmeans5 | 0.0039 | 0.3966 |
| kmeans4 | 0.0039 | 0.4017 |
| q5 | 0.0039 | 0.3791 |
| bikmeans3 | 0.0039 | 0.2693 |
| q4 | 0.0039 | 0.3962 |
| max25 | 0.0039 | 0.4664 |
| max50 | 0.0039 | 0.4744 |
| max75 | 0.0039 | 0.4827 |
| kmeans3 | 0.0039 | 0.4028 |
| q3 | 0.0039 | 0.388 |

**Table S6:** P-values and Mean Area Between the Curves (MABC) values of Different Discretization for DiscreeTest, Hepatocytic Cell Signaling Network

| Discretization | p-values | MABC |
|---|---|---|
| bikmeans2 | 0.059 | 0.137 |
| mean | p<0.001 | 0.07 |
| TDT | p<0.001 | 0.074 |
| i2 | p<0.001 | 0.028 |
| q2 | p<0.001 | 0.067 |
| kmeans2 | 0.2503 | 0.14 |
| max25 | p<0.001 | 0.064 |
| max50 | p<0.001 | 0.066 |
| max75 | p<0.001 | 0.074 |
| top75 | 0.043 | 0.169 |
| top25 | p<0.001 | 0.303 |

thus intrinsically the datasets contain noise. We showed that our evaluation metric identifies an optimal discretization on both networks using their corresponding timeseries datasets.

**Assessment of noise impact before reverse-engineering methods are been applied:** The objective is to assess whether DiscreeTest can capture dynamic trends in the data, with different levels of noise.

We considered the *in silico* Pandapas network and its corresponding time-series datasets. Under $0\%, 1\%$ and $5\%$ noise levels in the time-series datasets, we compared the original time series and the discretized time series based on the number of local minima and local maxima. The presence of a local maximum portrays

**Table S7:** P-values and Mean Area Between the Curves (MABC) values of Different Discretization for DiscreeTest, IRMA Network

| Discretization | switch off | | switch on | |
|---|---|---|---|---|
| | p-values | MABC | p-values | MABC |
| bikmeans2 | p<0.001 | 0.1566 | 0.0042 | 0.466 |
| bikmeans3 | 0.0002 | 0.209 | 0.0042 | 0.441 |
| bikmeans4 | p<0.001 | 0.1889 | 0.0042 | 0.3881 |
| bikmeans5 | p<0.001 | 0.1817 | 0.0042 | 0.3961 |
| q4 | 0.0266 | 0.4824 | 0.0768 | 0.4617 |
| TDT | p<0.001 | 0.0497 | 0.0042 | 0.43 |
| i4 | 0.0002 | 0.1582 | 0.0042 | 0.3546 |
| i5 | 0.0002 | 0.1747 | 0.0042 | 0.3463 |
| q5 | 0.0072 | 0.4745 | 0.0213 | 0.4949 |
| kmeans2 | p<0.001 | 0.1405 | 0.0042 | 0.4178 |
| kmeans3 | 0.0002 | 0.1792 | 0.0042 | 0.3774 |
| kmeans4 | 0.0002 | 0.1797 | 0.0042 | 0.3681 |
| kmeans5 | 0.0002 | 0.1829 | 0.0042 | 0.3338 |
| mean | 0.0002 | 0.3311 | 0.0213 | 0.3743 |
| q3 | 0.1892 | 0.4624 | 0.2101 | 0.4941 |
| i2 | p<0.001 | 0.0972 | 0.0042 | 0.4053 |
| i3 | 0.0002 | 0.1436 | 0.0042 | 0.3505 |
| q2 | 1 | 0.4887 | 0.8036 | 0.4601 |

the trend of a time-series shifting from increasing to decreasing. A local minimum portrays the trend shifting from decreasing to increasing. We define a true positive as a local maximum that is present in both original time series and the discretized time series, a true negative as a local minimum that is present in both, the original time series and the discretized time series. A false positive is defined as a local maximum present only in the discretized data or a local minimum that is missed by the discretized data. A false negative is defined as a local minimum present only in the discretized data or a local maximum missed by the discretized data. We focus on discretization methods that either pass the qualification step (i2, top75, top25, mean) or gives small MABC (bikmeans2, bikmeans3, bikmeans4, bikmeans5) (shown in Table S5). We compute confusion matrices for different the discretization methods with different levels of noise (Table S8-S10). We show that the discretization methods identified by DiscreeTest (marked on yellow on Tables S8, S9 and S10 for data with $0\%, 1\%$ and $5\%$ noise levels, respectively) consistently give higher positive predictive value and high accuracy (marked on blue on Tables S8, S9 and S10 for data with $0\%, 1\%$ and $5\%$ noise levels, respectively).

We further provide a couple of visual examples. We show one time series from data with no noise (panel A, Figure S10), and two discretizations of it. The original time series (solid black line in panel A, Figure S10) has 1 local minimum (marked out by black square) and has no local maxima. The data discretized by i2, the discretization identified by DiscreeTest, captures the local minimum (marked on red hexagram). Another two-level discretization method, top75, overfits the dynamical trend and have 2 local maxima and 2 local minima (marked on yellow dots). Similarly, when comparing binary discretizations to multi-level discretization (panel B in Figure S10), the original time-series data shows a continuous decreasing, with no local minima nor local maxima. All the discretization methods (i2, mean, bikmeans5 discretizations) show a decreasing trend, however, bikmeans5 generates 2 local maxima and 2 local minima, much more than what is captured in i2 or mean discretization methods. This is also observed in the discretized data with 1% noise (panel C, Figure S10) and data with 5% noise (panel D, Figure S10). These examples show that the discretization method identified by DiscreeTest is optimal in comparison to the other discretization methods benchmarked.

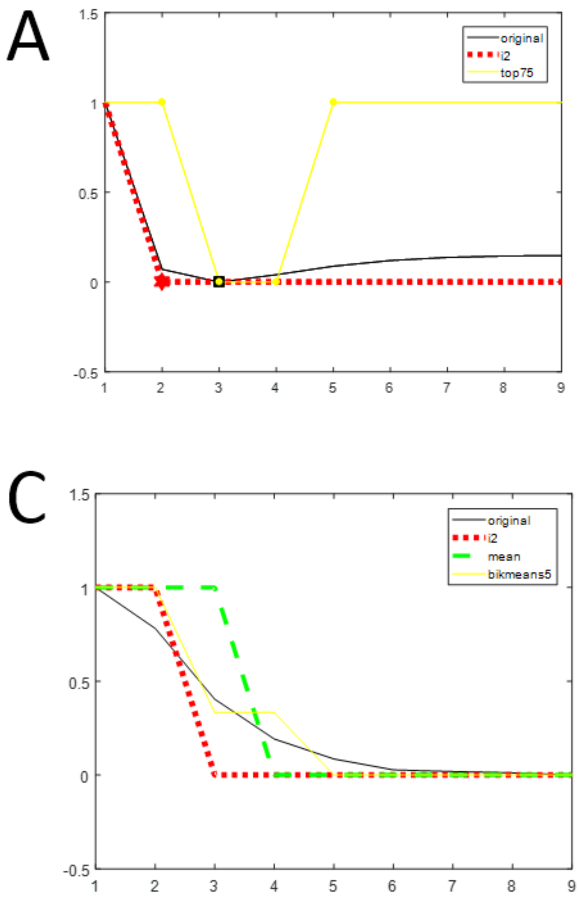


**Figure S10:** Examples from Pandapas network showing how well the discretized data capture the trend change in original data. These examples use the data from gene 2, time series 2 of Pandapas network. Examples span noiseless data (panel A and B), 1% noise data (panel C) and 5% noise data (panel D). In panel A, we show how binary discretized data (by either i2 or top75, in red and in yellow respectively) may catch the change of trend in the original data (black solid line) differently. Points of transitions (local maxima or local minima) are marked out using red hexagram in i2 discretization, black square in original data, and yellow dots in top75 discretized data. In panel B, we show how discretization with different levels (i2 in red dotted line, mean in green dash line, bikmeans5 in yellow solid line) may catch the change of trend in the original noiseless data (black solid line) differently. In panel C, we show how discretization with different levels (i2 in red dotted line, mean in green dash line, bikmeans5 in yellow solid line) may catch the change of trend in the original 1% noise data (black solid line) differently. In panel D, we show how discretization with different levels (i2 in red dotted line, mean in green dash line, bikmeans5 in yellow solid line) may catch the change of trend in the original 5% noise data (black solid line) differently.

**Table S8:** True positive (TP), false negative (FN), false positive (FP), true negative(TN), positive predictive value (PPV) and accuracy of a discretized data that captures dynamic changing patterns (local maxima or local minima) in the original data, noiseless Pandapas data

| Discretization | TP | FN | FP | TN | PPV | accuracy |
|---|---|---|---|---|---|---|
| i2 | 24 | 42 | 36 | 29 | 0.4 | 0.405 |
| top75 | 31 | 47 | 57 | 33 | 0.352 | 0.381 |
| top25 | 31 | 79 | 65 | 33 | 0.352 | 0.308 |
| mean | 27 | 56 | 0 | 33 | 1 | 0.517 |
| bikmeans5 | 24 | 31 | 49 | 16 | 0.328 | 0.333 |
| bikmeans4 | 24 | 23 | 41 | 16 | 0.369 | 0.384 |
| bikmeans3 | 24 | 23 | 41 | 16 | 0.369 | 0.384 |
| bikmeans2 | 18 | 23 | 28 | 16 | 0.391 | 0.4 |

**Table S9:** True positive (TP), false negative (FN), false positive (FP), true negative(TN), positive predictive value (PPV) and accuracy (ACC) of a discretized data that captures dynamic changing patterns (local maxima or local minima) in the original data, 1% noise Pandapas data

| Discretization | TP | FN | FP | TN | PPV | ACC |
|---|---|---|---|---|---|---|
| i2 | 24 | 23 | 41 | 16 | 0.369 | 0.384 |
| top75 | 31 | 55 | 65 | 33 | 0.322 | 0.347 |
| top25 | 31 | 95 | 81 | 33 | 0.276 | 0.266 |
| mean | 24 | 42 | 36 | 29 | 0.4 | 0.405 |
| bikmeans5 | 24 | 31 | 49 | 16 | 0.328 | 0.333 |
| bikmeans4 | 24 | 23 | 41 | 16 | 0.369 | 0.384 |
| bikmeans3 | 24 | 23 | 41 | 16 | 0.369 | 0.384 |
| bikmeans2 | 20 | 17 | 51 | 2 | 0.282 | 0.244 |

**Table S10:** True positive (TP), false negative (FN), false positive (FP), true negative(TN), positive predictive value (PPV) and accuracy (ACC) of a discretized data that captures dynamic changing patterns (local maxima or local minima) in the original data, 5% noise Pandapas data

| Discretization | TP | FN | FP | TN | PPV | ACC |
|---|---|---|---|---|---|---|
| i2 | 24 | 23 | 41 | 16 | 0.369 | 0.384 |
| top75 | 31 | 55 | 65 | 33 | 0.322 | 0.347 |
| top25 | 31 | 95 | 81 | 33 | 0.276 | 0.266 |
| mean | 24 | 42 | 36 | 29 | 0.4 | 0.405 |
| bikmeans5 | 24 | 31 | 49 | 16 | 0.328 | 0.333 |
| bikmeans4 | 24 | 23 | 41 | 16 | 0.369 | 0.384 |
| bikmeans3 | 24 | 23 | 41 | 16 | 0.369 | 0.384 |
| bikmeans2 | 20 | 17 | 51 | 2 | 0.282 | 0.244 |

### References


D. Camacho, P. Vera-Licona, P. Mendes, and R. Laubenbacher. Comparison of reverse-engineering methods using an in silico network. *Annals of the New York Academy of Sciences*, 1115(1):73–89, 2007.

I. Cantone, L. Marucci, F. Iorio, M. A. Ricci, V. Belcastro, M. Bansal, S. Santini, M. Di Bernardo, D. Di Bernardo, and M. P. Cosma. A yeast synthetic network for in vivo assessment of reverse-engineering and modeling approaches. *Cell*, 137(1):172–181, 2009.

S. Erdal, O. Ozturk, D. Armbruster, H. Ferhatosmanoglu, and W. C. Ray. A time series analysis of microarray data. In *Bioinformatics and Bioengineering, 2004. BIBE 2004. Proceedings. Fourth IEEE Symposium on*, pages 366–375. IEEE, 2004.

C. A. Gallo, J. A. Carballido, and I. Ponzoni. Discovering time-lagged rules from microarray data using gene profile classifiers. *BMC bioinformatics*, 12(1):1, 2011.

C. A. Gallo, R. L. Cecchini, J. A. Carballido, S. Micheletto, and I. Ponzoni. Discretization of gene expression data revised. *Briefings in Bioinformatics*, 17(5):758–770, 2016. doi: 10.1093/bib/bbv074.

S. Gaudet, K. A. Janes, J. G. Albeck, E. A. Pace, D. A. Lauffenburger, and P. K. Sorger. A compendium of signals and responses triggered by prodeath and prosurvival cytokines. *Molecular & Cellular Proteomics*, 4(10):1569–1590, 2005.

L. Ji and K.-L. Tan. Mining gene expression data for positive and negative co-regulated gene clusters. *Bioinformatics*, 20(16):2711–2718, 2004.

S. Klamt, J. Saez-Rodriguez, J. A. Lindquist, L. Simeoni, and E. D. Gilles. A methodology for the structural and functional analysis of signaling and regulatory networks. *BMC bioinformatics*, 7(1):1, 2006.

P. Li, P. Gong, H. Li, E. J. Perkins, N. Wang, and C. Zhang. Gene regulatory network inference and validation using relative change ratio analysis and time-delayed dynamic bayesian network. *EURASIP Journal on Bioinformatics and Systems Biology*, 2014(1):1, 2014.

Y. Li, L. Liu, X. Bai, H. Cai, W. Ji, D. Guo, and Y. Zhu. Comparative study of discretization methods of microarray data for inferring transcriptional regulatory networks. *BMC bioinformatics*, 11(1):520, 2010.

S. C. Madeira and A. L. Oliveira. An evaluation of discretization methods for non-supervised analysis of time-series gene expression data. *Instituto de Engenharia de Sistemas e Computadores Investigacao e Desenvolvimento, Technical Report*, 42, 2005.

D. Marbach, T. Schaffter, C. Mattiussi, and D. Floreano. Generating realistic in silico gene networks for performance assessment of reverse engineering methods. *Journal of computational biology*, 16(2):229–239, 2009.

D. Marbach, R. J. Prill, T. Schaffter, C. Mattiussi, D. Floreano, and G. Stolovitzky. Revealing strengths and weaknesses of methods for gene network inference. *Proceedings of the national academy of sciences*, 107(14):6286–6291, 2010.

C. S. Möller-Levet, K. Chu, and O. Wolkenhauer. Dna microarray data clustering based on temporal variation: Fcv with tsd preclustering. *Applied Bioinformatics*, 2(1):35–45, 2003.

M. K. Morris, J. Saez-Rodriguez, D. C. Clarke, P. K. Sorger, and D. A. Lauffenburger. Training signaling pathway maps to biochemical data with constrained fuzzy logic: quantitative analysis of liver cell responses to inflammatory stimuli. *PLoS Comput Biol*, 7(3): e1001099, 2011.

M. K. Morris, I. Melas, and J. Saez-Rodriguez. Construction of cell type-specific logic models of signaling networks using cellnopt. *Computational Toxicology: Volume II*, pages 179–214, 2013.

I. Ponzoni, F. Azuaje, J. Augusto, and D. Glass. Inferring adaptive regulation thresholds and association rules from gene expression data through combinatorial optimization learning. *IEEE/ACM Transactions on Computational Biology and Bioinformatics*, 4(4): 624–634, 2007.

R. J. Prill, D. Marbach, J. Saez-Rodriguez, P. K. Sorger, L. G. Alexopoulos, X. Xue, N. D. Clarke, G. Altan-Bonnet, and G. Stolovitzky. Towards a rigorous assessment of systems biology models: the dream3 challenges. *PloS one*, 5(2):e9202, 2010.

J. Saez-Rodriguez, L. Simeoni, J. A. Lindquist, R. Hemenway, U. Bommhardt, B. Arndt, U.-U. Haus, R. Weismantel, E. D. Gilles, S. Klamt, et al. A logical model provides insights into t cell receptor signaling. *PLoS Comput Biol*, 3(8):e163, 2007.

J. Saez-Rodriguez, A. Goldsipe, J. Muhlich, L. G. Alexopoulos, B. Millard, D. A. Lauffenburger, and P. K. Sorger. Flexible informatics for linking experimental data to mathematical models via datarail. *Bioinformatics*, 24(6):840–847, 2008.

J. Saez-Rodriguez, L. G. Alexopoulos, J. Epperlein, R. Samaga, D. A. Lauffenburger, S. Klamt, and P. K. Sorger. Discrete logic modelling as a means to link protein signalling networks with functional analysis of mammalian signal transduction. *Molecular systems biology*, 5(1):331, 2009.

L. A. Soinov, M. A. Krestyaninova, and A. Brazma. Towards reconstruction of gene networks from expression data by supervised learning. *Genome biology*, 4(1):1, 2003.

C. Terfve, T. Cokelaer, D. Henriques, A. MacNamara, E. Goncalves, M. K. Morris, M. van Iersel, D. A. Lauffenburger, and J. Saez-Rodriguez. Cellnoptr: a flexible toolkit to train protein signaling networks to data using multiple logic formalisms. *BMC systems biology*, 6(1):1, 2012.

J. Yu, V. A. Smith, P. P. Wang, A. J. Hartemink, and E. D. Jarvis. Advances to bayesian network inference for generating causal networks from observational biological data. *Bioinformatics*, 20(18):3594–3603, 2004.

M. Zou and S. D. Conzen. A new dynamic bayesian network (dbn) approach for identifying gene regulatory networks from time course microarray data. *Bioinformatics*, 21(1):71–79, 2005.
